# Identification and dynamics of the DHHC16-DHHC6 palmitoylation cascade

**DOI:** 10.1101/134007

**Authors:** Laurence Abrami, Tiziano Dallavilla, Patrick A. Sandoz, Mustafa Demir, Béatrice Kunz, Georgios Savoglidis, Vassily Hatzimanikatis, F. Gisou van der Goot

**Author notes:** Contributed equally to this work (Laurence Abrami, Tiziano Dallavilla).

## Abstract

S-Palmitoylation is the only reversible post-translational lipid modification. Knowledge about the DHHC family of palmitoyltransferases is very limited. Here we show that mammalian DHHC6, which modifies key proteins of the endoplasmic reticulum, is controlled by an upstream palmitoyltransferase, DHHC16, revealing the first palmitoylation cascade. Combination of site specific mutagenesis of the three DHHC6 palmitoylation sites, experimental determination of kinetic parameters and data-driven mathematical modelling allowed us to obtain detailed information on the 8 differentially palmitoylated DHHC6 species. We found that species rapidly interconvert through the action of DHHC16 and the Acyl Protein Thioesterase APT2, that each species varies in terms of turnover rate and activity, altogether allowing the cell to robustly tune its DHHC6 activity.

## INTRODUCTION

Cells constantly interact and respond to their environment. This requires tight control of protein function in time and in space, which largely occurs through reversible post-translational modifications of proteins, such as phosphorylation, ubiquitination and S-palmitoylation. The latter consist in the addition on an acyl chain, generally C16 in mammals, to cysteine residues, thereby altering the hydrophobicity of the protein and tuning its function (Blaskovic et al, 2014; Chamberlain & Shipston, 2015). More precisely, palmitoylation may controls the interaction of a proteins with membranes or specific membrane domains, affect its conformation, trafficking, stability and/or activity (Blaskovic et al, 2014; Chamberlain et al, 2013; Conibear & Davis, 2010; Fukata et al, 2016). In the cytosol, the acyl chain is attached to the protein via a thioester bond through the action of DHHC palmitoyltransferases, a family of multispanning transmembrane proteins (Blaskovic et al, 2014; Chamberlain et al, 2013; Conibear & Davis, 2010; Fukata et al, 2016).

The list of proteins undergoing palmitoylation is ever increasing ((Blanc et al, 2015), http://swisspalm.epfl.ch/) and the modification is found to be important in numerous key cellular processes including neuronal development and activity (Fukata & Fukata, 2010), cardiac function (Pei et al, 2016), systemic inflammation (Beard et al, 2016), innate immunity to viruses (Mukai et al, 2016), cell polarity (Chen et al, 2016), EGF-signalling (Runkle et al, 2016), protease activity (Skotte et al, 2017) and cancer (Coleman et al, 2016; Thuma et al, 2016).

While novel roles and targets of palmitoylation are constantly reported, little is known about the regulation and dynamics of this post-translational modification. Here we focused on one of the 23 human palmitoyltransferase, DHHC6, which localizes to the endoplasmic reticulum (ER) and controls a panel of key ER substrates such as the ER chaperone calnexin (Lakkaraju et al, 2012), the ER E3 ligase gp78 (Fairbank et al, 2012), the IP3 receptor (Fredericks et al, 2014) as well as cell surface proteins such as the transferrin receptor (Senyilmaz et al, 2015). For each of these proteins, palmitoylation controls stability, localization, trafficking and/or function.

As most DHHC enzymes, DHHC6 is a tetra-spanning membrane protein. It has a short N-terminal extension and a long C-terminal tail composed of an approx. 100 residue domain of unknown structure followed by a variant SH3_2 domain (Fig. 1A, Fig.1-figure supplement 1). At the C-terminus, it contains a KKNR motif which when transferred to the DHHC3 enzyme leads to its relocalization from the Golgi to the ER (Gorleku et al, 2011), suggesting it contributes to the DHHC6 ER localization.

**Figure 1.**
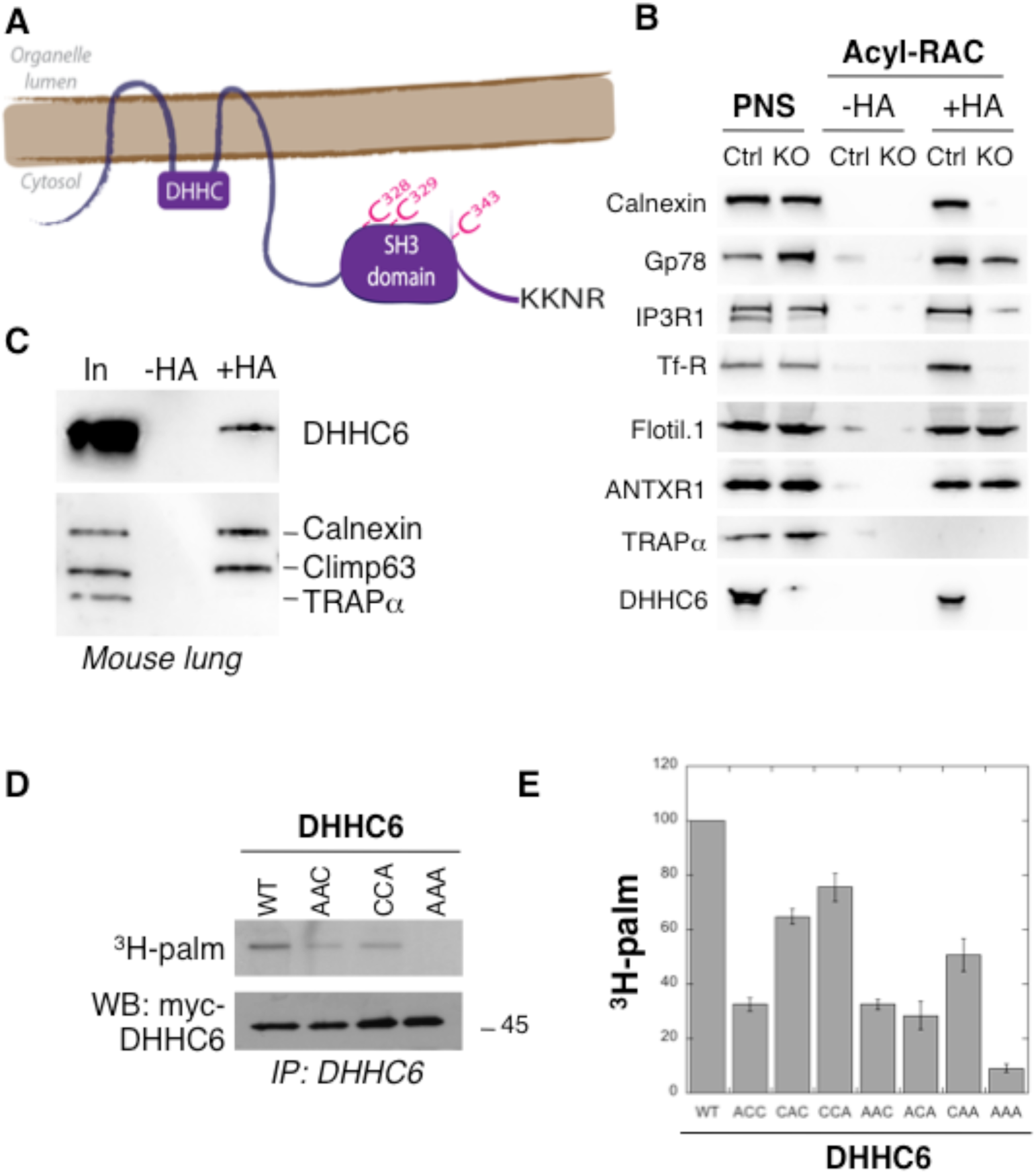
DHHC6 can undergo palmitoylation on three cysteines of its SH3 domain. **A.** Schematic representation of DHHC6 enzyme. The potential ER retention motif is KKNR. **B.** Analysis of protein acylation in control HAP cells (Ctrl) versus DHHC6 KO HAP (KO). HAP cell membranes were recovered by centrifugation and incubated with MMTS and then with hydroxylamine (+HA) or with Tris (-HA) together with free thiol group binding beads. Eluted fractions were analyzed by immunoblotting with the indicated antibodies. PNS represents 1/10 of the input fraction. **C.** DHHC6 acylation in mouse tissues. 400µg total proteins extracted from mouse lung were incubated with MMTS and then with hydroxylamine (+HA) or with Tris (-HA) together with free thiol group binding beads. Eluted fractions were analyzed by immunoblotting with the indicated antibodies. “In” represents 1/10 of the input fraction. Calnexin and the ER shaping protein CLIMP-63 (Lakkaraju et al, 2012; Schweizer et al, 1993) were used as positive controls and the cysteine-less protein Trapα as negative control. **D.** Palmitoylation of DHHC6 cysteine mutants. Hela cells were transfected with plasmids encoding WT or the indicated Myc-tagged DHHC6 mutant constructs for 24h. Cells were then metabolically labelled for 2 hours at 37°C with ^3^H-palmitic acid. Proteins were extracted, immunoprecipitated with Myc antibodies, subjected to SDS-PAGE and analysed by autoradiography (^3^H-palm), quantified using the Typhoon Imager or by immunoblotting with the indicated antibodies. **E.** Quantification of ^3^H-palmitic acid incorporation into DHHC6. Quantified values were normalized to protein expression level. The calculated value of ^3^H-palmitic acid incorporation into WT DHHC6 was set to 100% and all mutants were expressed relative to this (n=4, error bars represent standard deviation).

Here we show that DHHC6 function, localization and stability are all regulated through the dynamic multi-site palmitoylation of its SH3_2 domain. Interestingly, palmitoylation occurs via an upstream palmitoyltransferase, DHHC16, revealing for the first time the existence of palmitoylation cascades. Depalmitoylation is mediated by the Acyl Protein Thioesterase APT2. Palmitoylation can occur on three different sites, leading to potentially 8 different DHHC6 species defined by acyl site occupancy that can interconvert through the actions of DHHC16 and APT2. Using mathematical modelling combined with various kinetic measurements on WT and mutant DHHC6, we probed the complexity of this system and its importance for DHHC6 function. Altogether, this study shows that palmitoylation affects the quaternary assembly of DHHC6, its localization, stability and function. Moreover we show that the presence of 3 sites is necessary for the robust control of DHHC6 activity.

## RESULTS

### Palmitoylation of the DHHC6 SH3_2 domain

DHHC6 KO cells were generated using the CRISPR-cas9 system in the near haploid cell line HAP1. Using the Acyl-RAC capture method to isolate palmitoylated proteins (Werno & Chamberlain, 2015), we verified that the ER chaperone calnexin (Lakkaraju et al, 2012), the E3 ligase gp78 (Fairbank et al, 2012), the IP3 receptor (Fredericks et al, 2014) and the transferrin receptor (Senyilmaz et al, 2015) are indeed DHHC6 targets (Fig. 1B). Interestingly calnexin and the transferrin receptor were no longer captured by Acyl-RAC in the HAP1 DHHC6 KO cells, confirming that they are exclusively modified by DHHC6, while capture of the IP3 receptor and gp78 was reduced but not abolished indicating that they can also be modified by other palmitoyltransferases (Fig. 1B) consistent with previous findings (Fairbank et al, 2012; Fredericks et al, 2014). As negative controls, we tested flotillin1, a known target of DHHC5 (Li et al, 2012) and the anthrax toxin receptor 1 modified by a yet to be determined DHHC enzyme (Abrami et al, 2006).

When probing samples for DHHC6 itself, we found that the palmitoyltransferase itself was captured by Acyl-RAC in HAP1 cells (Fig. 1B) and also in different mouse tissues (shown for mouse lung tissue in Fig. 1C). Palmitoylation of DHHC6 was reported in a large-scale proteomics analysis to occur on Cys-328, Cys-329 and Cys-343 in the SH3 domain (Yang et al, 2010). These cysteines are conserved in vertebrates (Fig.1-figure supplement 1). We generated single (ACC, CAC, CCA), double (AAC, CAA, ACA) and triple (AAA) cysteine-to-alanine mutants and monitored ^3^H-palmitate incorporation during 2hrs. Palmitoylation of DHHC6 WT, but not the AAA mutant, was readily detected (Fig. 1DE). All single cysteine mutants, especially C328A, and all double mutants showed a decrease in the ^3^H-palmitate signal (Fig. 1DE). Thus DHHC6 can indeed undergo palmitoylation, and importantly it has no palmitoylation sites other than the 3 in the SH3 domain.

### The DHHC16-DHHC6 palmitoylation cascade

We next investigated whether DHHC6 was palmitoylating itself, in *cis* or *trans*. We generated an inactive DHHC deletion mutant (ΔDHHC) as well as a cell line stably expressing an shRNA against DHHC6 (Fig. 2-figure supplement 1A). WT and ΔDHHC DHHC6 both underwent palmitoylation when transiently expressed in control or shRNA DHHC6 cells (Fig. 2-figure supplement 1A), indicating that DHHC6 must be modified by another enzyme. We next performed ^3^H-palmitate labelling experiments upon siRNA silencing of each of the 23 DHHC enzymes. Only silencing of DHHC16 led to a major, 60%, drop in signal intensity (Fig. 2A). ^3^H-palmitate incorporation was also reduced for the AAC and CCA mutants (Fig. 2-figure supplement 1B), showing that all three sites are modified by the same enzyme.

**Figure 2.**
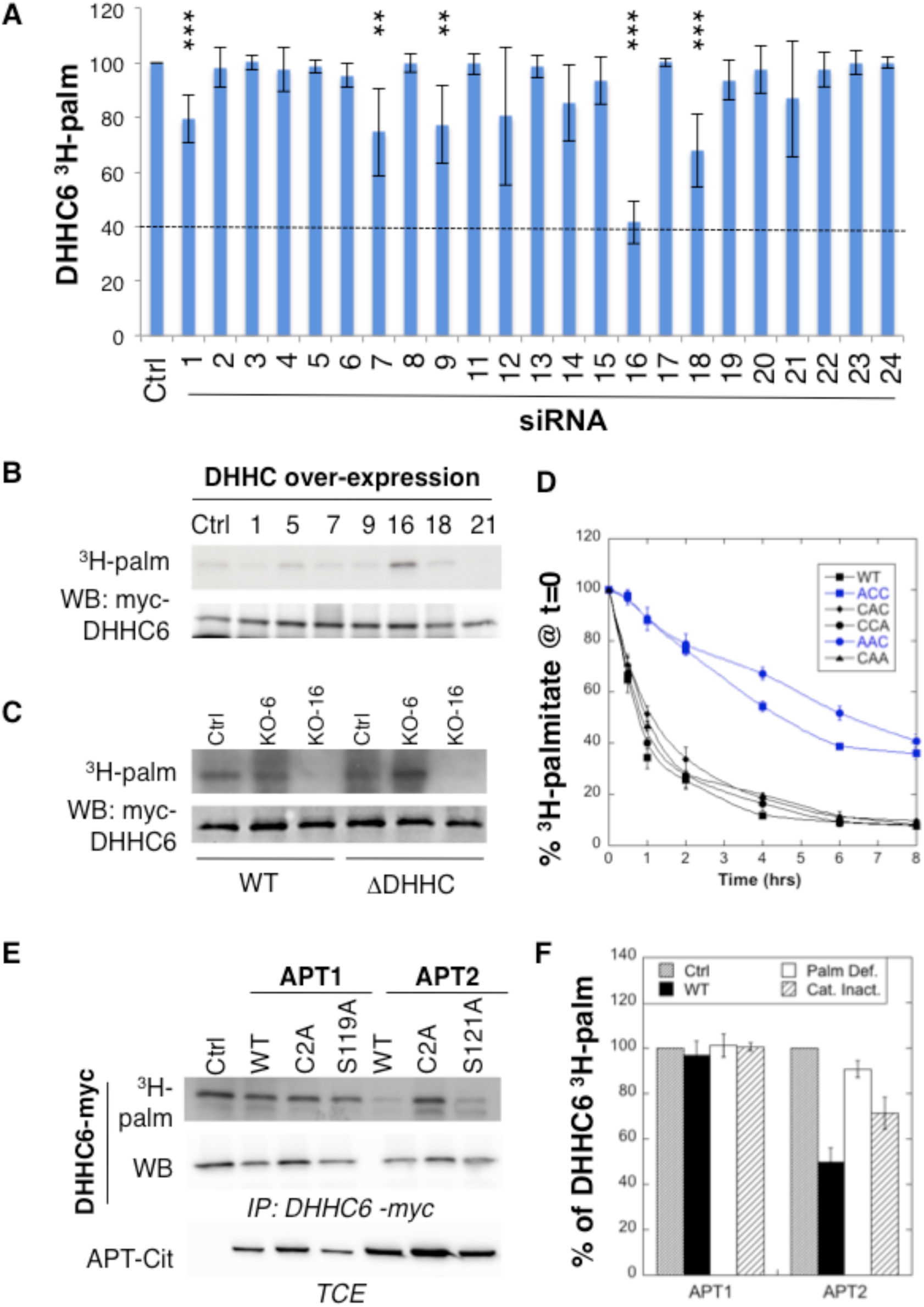
DHHC6 is palmitoylated by DHHC16 and depalmitoylated by APT2. **A.** Identification of the DHHC6 palmitoyltransferase by siRNA screening of DHHC enzymes. Hela cells were transfected with siRNAs against the indicated DHHC enzyme for 72h and with the myc-tagged WT DHHC6 expressing construct for the last 24h. Cells were then metabolically labelled 2 hours at 37°C with ^3^H-palmitic acid. Proteins were extracted, immunoprecipitated with myc antibodies and subjected to SDS-PAGE and analysed by autoradiography, quantified using the Typhoon Imager or by immunoblotting with myc antibodies. ^3^H-palmitic acid incorporation into DHHC6 was quantified and normalized to protein expression levels. The calculated value of ^3^H-palmitic acid incorporation into DHHC6 was set to 100% for an irrelevant siRNA (Ctrl) and all siRNA were expressed relative to this (n=6, error bars represent standard deviation). **B.** Identification of the DHHC6 palmitoyltransferase by DHHC over-expression. Hela cells were transfected with indicated the DHHC constructs and with myc-tagged WT DHHC6 construct for 24h. Cells were then metabolically labelled 2 hours at 37°C with ^3^H-palmitic acid. Proteins were extracted, immunoprecipitated with myc antibodies and subjected to SDS-PAGE and analysed by autoradiography (^3^H-palm) or by immunoblotting with myc antibodies. **C.** Analysis of DHHC6 acylation in control HAP cells (Ctrl) versus DHHC6 KO HAP (KO-6) or DHHC16 KO HAP (KO-16). Cells were transfected with the myc-tagged WT DHHC6 construct for 24h, then metabolically labelled 2 hours at 37°C with ^3^H-palmitic acid. Proteins were extracted, immunoprecipitated with myc antibodies and subjected to SDS-PAGE and analysed by autoradiography (^3^H-palm) or by immunoblotting with myc antibodies. **D.** Palmitoylation decay of WT or mutant DHHC6. Hela cells were transfected with plasmids encoding WT or the indicated mutant Myc-tagged DHHC6 constructs for 24h. Cells were then metabolically labelled 2 hours at 37°C with ^3^H-palmitic acid, washed and incubated with complete medium for different hours. Proteins were extracted, immunoprecipitated with myc antibodies and subjected to SDS-PAGE and analysed by autoradiography, quantified using the Typhoon Imager or by immunoblotting with anti-myc antibodies. ^3^H-palmitic acid incorporation was quantified for each time point, normalized to protein expression level. ^3^H-palmitic acid incorporation was set to 100% at t=0 after the 2 hours pulse and all different times of chase were expressed relative to this (n=4, error bars represent standard deviation). **E.** DHHC6 palmitoylation upon APT overexpression. Hela cells were transfected with plasmids encoding myc-tagged WT DHHC6 and the indicated mutant citrin-tagged APT1 or APT2 constructs for 24h. Cells were then metabolically labelled 2 hours at 37°C with ^3^H-palmitic acid. Proteins were extracted, immunoprecipitated with myc antibodies and subjected to SDS-PAGE and analysed by autoradiography (^3^H-palm), quantified using the Typhoon Imager or by immunoblotting with anti-myc antibodies. **F.** Quantification of ^3^H-palmitic acid incorporation into DHHC6. Quantified values were normalized to protein expression level. ^3^H-palmitic acid incorporation was set to 100% for control cells (Ctrl) and values obtained for APT overexpressing cells were expressed relative to this (n=3, error bars represent standard deviation).

A screen by over-expression of DHHC enzymes also pointed to DHHC16 as the responsible enzyme (Fig. 2B, Fig. 2-figure supplement 1C). The final confirmation was obtained using CRISPR-Cas9-generated DHHC16 KO HAP1 cells. WT and ΔDHHC DHHC6 both incorporated ^3^H-palmitate when expressed in the control and the DHHC6 KO cells, but not in DHHC16 KO cells (Fig. 2C). Consistent with these findings, co-immunoprecipitation experiments following transient over-expression of myc-tagged DHHC6 and FLAG-tagged DHHC16 showed that the two enzymes can interact (Fig. 2-figure supplement 1D). Interestingly, *zdhhc6* silencing in Hela cells or *zdhhc6* knock out in HAP1 cells both led to an increase in the *zdhhc16* mRNA (Fig. 2-figure supplement 2AB), indicating that these two proteins interact physically and genetically.

Altogether these experiments show that DHHC6 can be palmitoylated on all three of its SH3_2 cysteine residues by DHHC16, revealing for the first time that palmitoylation can occur in a cascade, as occurs for phosphorylation for example in the MAK kinase pathway. DHHC16, which localizes under over-expression conditions to the ER and the Golgi (Fig. 2-figure supplement 3A), is not palmitoylated itself, as shown both by ^3^H-palmitate incorporation and Acyl-RAC (Fig. 2-figure supplement 2BC).

### Rapid APT2-mediated DHHC6 depalmitoylation

Palmitoylation is a reversible modification and thus has the potential to be dynamic. To analyse palmitate turnover on DHHC6, we performed ^3^H-palmitate pulse-chase experiments. Following a 2hr pulse, 50% of the ^3^H-palmitate was lost from DHHC6 in ≈ 1 hour, indicating rapid turnover of the acyl chains (Fig. 2D). We performed similar experiments on the various single and double cysteine mutants and found that rapid turnover required the presence of Cys-328. In its absence, the apparent palmitate release rates were considerably slower, reaching more than 4 hrs (Fig. 2D).

Depalmitoylation is an enzymatic reaction that is mediated by poorly characterized Acyl Protein Thioesterases (APT). We tested the involvement of APT1 and APT2 (Blaskovic et al, 2014), which are themselves palmitoylated on Cys-2 (Kong et al, 2013). We generated palmitoylation deficient variants of APT1 and 2 as well as catalytically inactive versions (APT1 S119A and APT2 S121A). Overexpression of WT or mutant APT1 had no detectable effect on DHHC6 palmitoylation (Fig. 2E, F). In contrast, overexpression of WT, but not palmitoylation deficient, APT2 led to a significant decrease in DHHC6 palmitoylation. APT2^S121A^ had an intermediate effect, possibly due to the formation of heterodimers between mutant and endogenous APT2 (Kong et al, 2013; Vujic et al, 2016). These observations show that DHHC6 palmitoylation is dynamic, in particular on Cys-328, and that depalmitoylation is mediated by APT2.

### Palmitoylation of Cys-328 destabilizes DHHC6

We next analysed the effect of palmitoylation on DHHC6 stability. Palmitoylation has been reported to affect protein turnover, increasing the half-life of proteins such as calnexin (Dallavilla et al, 2016) and the death receptor Fas (Rossin et al, 2015) but decreasing that of others such as gp78 (Fairbank et al, 2012). Protein turnover was monitored using ^35^S-Cys/Met metabolic pulse-chaseMyc-DHHC6 Hela cells, transiently expressing myc-DHHC6, were submitted to a 20 min metabolic pulse followed by different times of chase before lysis and anti-myc immunoprecipitation, SDS-PAGE and auto-radiography. Decay of newly synthesized DHHC6 was biphasic, with 40% undergoing gradual degradation during the first 5 hrs, and the remaining 60% undergoing degradation at a greatly reduced rate (Fig. 3A). The overall apparent half-life was 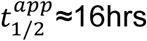. To enhance DHHC6 palmitoylation, we overexpressed DHHC16 or silenced APT2 expression. Both these genetic manipulations led to a dramatic acceleration of DHHC6 decay, with 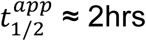 and 3hrs upon DHHC16 over expression and APT2 silencing, respectively (Fig. 3A). Remarkably, mutation of Cys-328, but not of Cys-329 or Cys-343, abolished the sensitivity to DHHC16 overexpression or APT2 silencing (Fig. 3BC, Fig. 3-figure supplement 1AB). APT2 silencing leads to ubiquitination of DHHC6 (Fig. 3D) and DHHC6 degradation can be rescued by the proteasome inhibitor MG132 (Fig. 3E). Thus altogether these observations indicate that palmitoylation of Cys-328 renders DHHC6 susceptible to degradation by the ERAD pathway.

**Figure 3.**
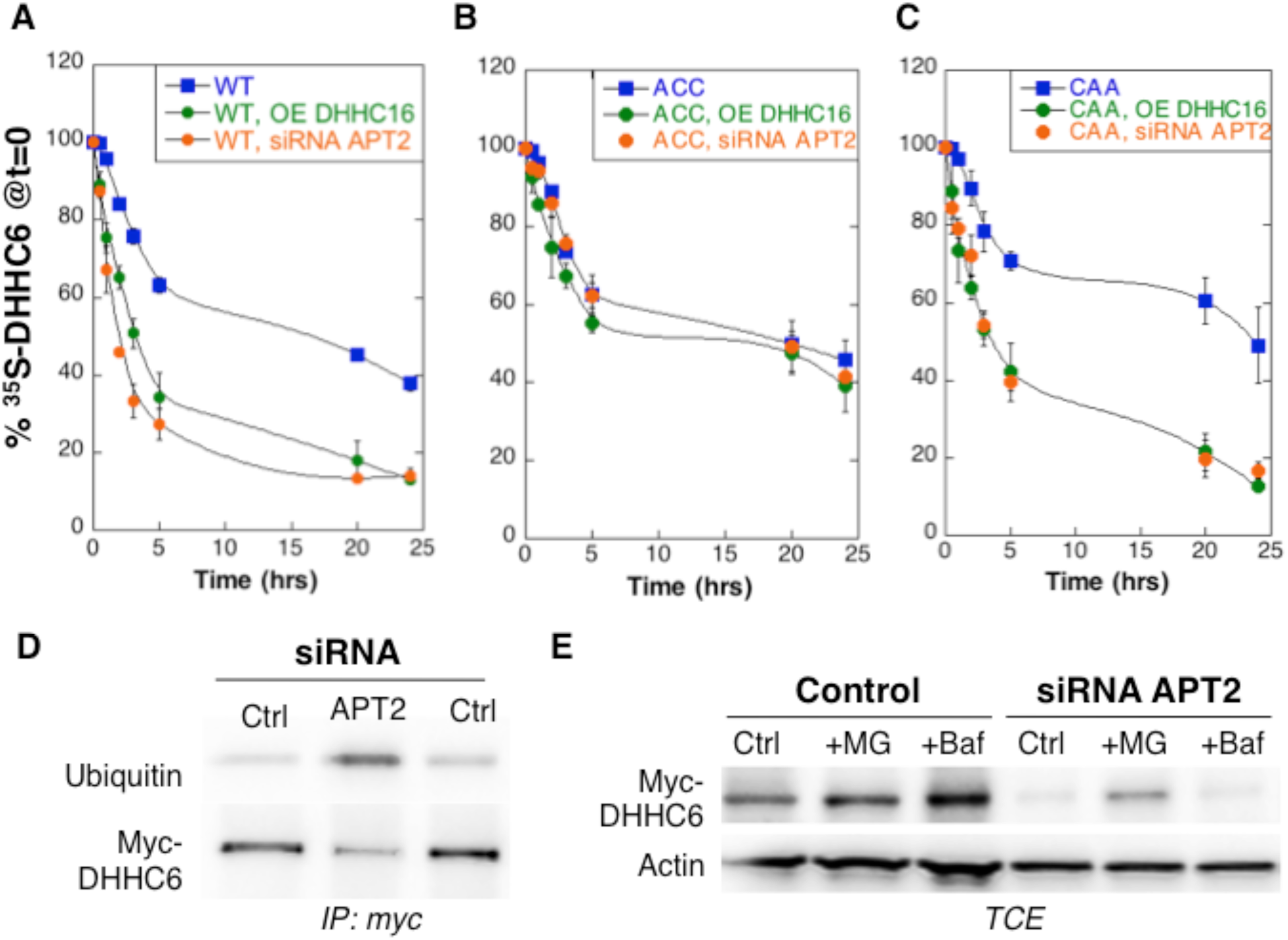
Palmitoylation on Cys-328 targets DHHC6 to ERAD. **ABC.** Degradation kinetics of DHHC6. Hela cells were transfected with plasmids encoding myc-tagged WT DHHC6 or cysteine mutants with or without FLAG-tagged WT DHHC16 for 24h after 48h transfection with siRNA APT2 or with control siRNA. Hela cells were incubated for 20min with ^35^S-Met/Cys at 37°C, washed and further incubated for different times at 37°C in complete medium. DHHC6 was immunoprecipitated and subjected to SDS-PAGE and analysed by autoradiography, quantified using the Typhoon Imager, and western blotting with anti-myc antibodies. ^35^S-Met/Cys incorporation was quantified for each time point, normalized to protein expression levels. ^35^S-Met/Cys incorporation was set to 100% for t=0 after the 20min pulse and all different times of chase were expressed relative to this (n=3, error bars represent standard deviation). **DE.** DHHC6 ubiquitination and proteasomal degradation. Hela cells were transfected with plasmids encoding myc-tagged WT DHHC6 constructs for 24h after 48h transfection with control (Ctrl) or APT2 siRNA. Proteins were extracted; DHHC6 was immunoprecipitated, subjected to SDS-PAGE and then analysed by immunoblotting with anti-ubiquitin or anti-myc antibodies (**D**). **E.** Cells were treated 4 hours with 10µM MG132 or with 100 nM Bafilomycin A before proteins were extracted; 40µg of total extract were subjected to SDS-PAGE and then analysed by immunoblotting against actin, used as equal loading control, or myc.

### Palmitoylation-dependent DHHC6 localization

We next investigated whether palmitoylation affects DHHC6 localization. WT DHHC6 shows a typical ER staining, co-localizing with Bip, a lumenal ER chaperone, and BAP31, a transmembrane ER protein (Fig. 4AB). Ectopically expressed WT DHHC6 sometimes also localized to a dot (Fig. 4AB). The DHHC6 AAA mutant was also clearly present in the reticular ER, but it tended to accumulate more frequently in dot-like ER structures, which stained positive for BAP31 but not BIP (Fig. 4AB). Thus the inability of DHHC6 AAA to undergo palmitoylation influences its localization within the two dimensional space of the ER membrane.

**Figure 4.**
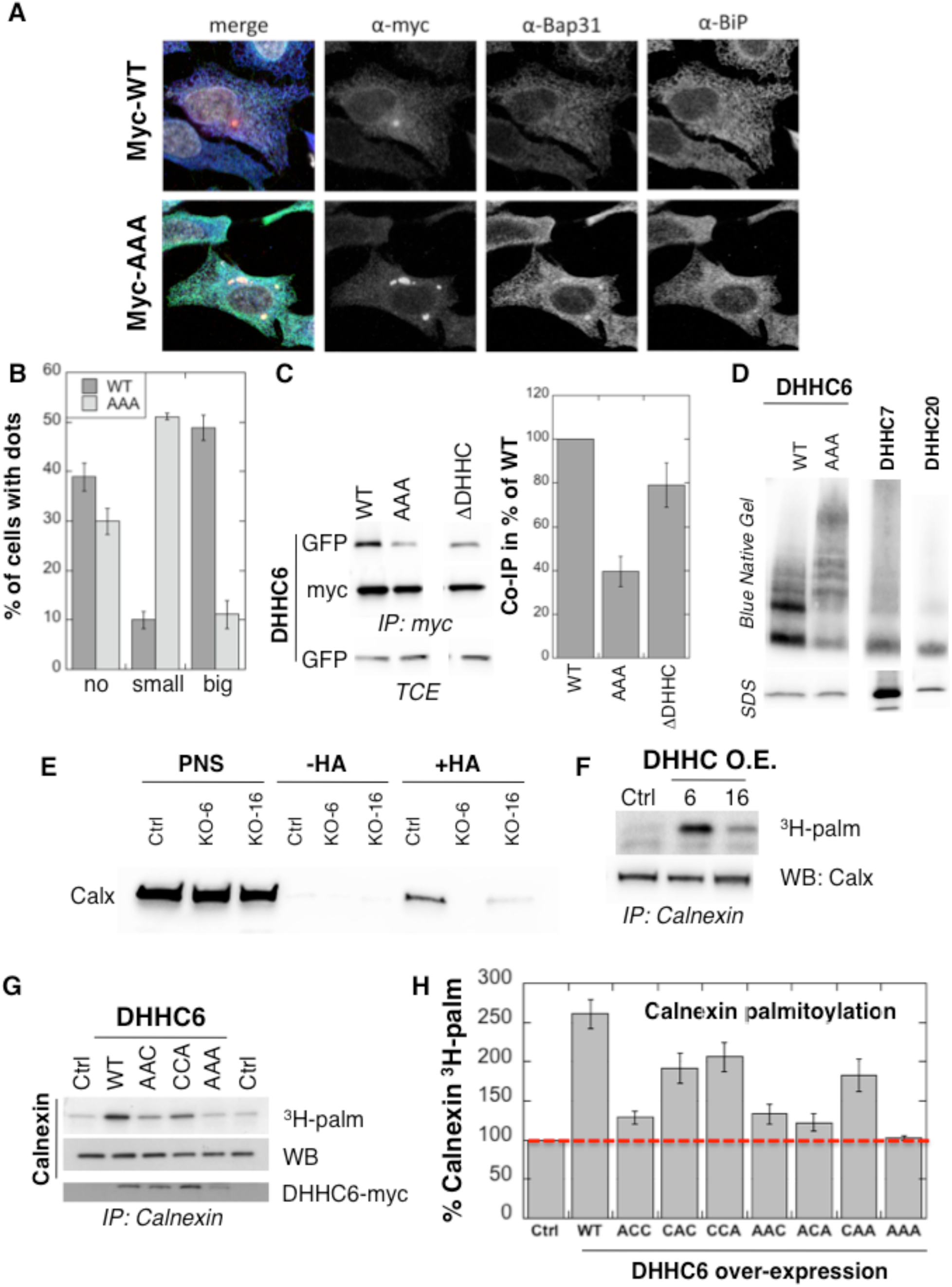
Pamitoylation affects assembly, localisation and function of DHHC6. **AB.** Immunofluorescence staining of HeLa cells transfected with myc-DHHC6 WT or AAA. The presence of dots (small or big) was reported for each cell and quantified. **C.** Co-immunoprecipitation of DHHC6 variants. Hela cells were transfected with plasmids encoding WT myc-DHHC6 and the indicated GFP-tagged mutant for 24h. Proteins were extracted, a total cell extract was analysed (TCE) and proteins were immunoprecipitated with myc antibodies and subjected to SDS-PAGE, then analysed by immunoblotting with anti-myc or anti-GFP antibodies. Quantification was performed by densitometry. The calculated value of co-immunoprecipitation with WT DHHC6 was set to 100%, all DHHC6 mutants were expressed relative to this (n=4, errors represent standard deviations). **D.** DHHC6 complexes. Hela cells were transfected with plasmids encoding myc-DHHC6 WT or mutant, or myc-tagged DHHC7 or DHHC20 for 24h. Proteins were extracted, 40µg of a TCE was analysed by SDS-PAGE or on blue native gels then analysed by immunoblotting with anti-myc antibodies. **E.** Analysis of endogenous calnexin acylation in control HAP cells (Ctrl) versus HAP cells KO for DHHC6 (KO-6) or for DHHC16 (KO-16). HAP cell membranes were recovered by centrifugation and incubated with MMTS and then with hydroxylamine (+HA) or with Tris (-HA) together with free thiol group binding beads. Eluted fractions were analysed by immunoblotting with anti-calnexin antibodies. PNS represents 1/10 of the input fraction. **F.** Analysis of endogenous calnexin palmitoylation in Hela cells overexpressing DHHC6 or DHHC16. Hela cells were transfected with plasmids encoding WT myc-DHHC6 or WT myc-DHHC16 constructs for 24h. Cells were then metabolically labelled 2 hours at 37°C with ^3^H-palmitic acid. Proteins were extracted and immunoprecipitated with anti-calnexin antibodies, subjected to SDS-PAGE and analysed by autoradiography (^3^H-palm) or by immunoblotting with anti-calnexin antibodies. **GH.** Analysis of endogenous calnexin palmitoylation in Hela cells overexpressing DHHC6 mutants. Hela cells were transfected with control plasmid (Ctrl) or plasmids encoding WT or mutants myc-DHHC6 for 24h. Cells were then metabolically labelled 2 hours at 37°C with ^3^H-palmitic acid. Proteins were extracted and immunoprecipitated with anti-calnexin antibodies, subjected to SDS-PAGE and analysed by autoradiography (^3^H-palm), quantified using the Typhoon Imager or by immunoblotting with anti-calnexin or anti-myc antibodies. ^3^H-palmitic acid incorporation into calnexin was set to 100% for the control plasmid and all DHHC6 mutants were expressed relative to this (n=4, errors represent standard deviations).

Certain DHHC enzymes were reported to dimerize (Lai & Linder, 2013). We therefore tested whether DHHC6 also dimerizes, and if so whether assembly is influenced by palmitoylation. Immunoprecipitation experiments using DHHC6 constructs with two different tags showed that WT DHHC6 can associate with itself as well as with the ΔDHHC variant but not with the AAA mutant (Fig. 4C). Blue native gel analysis confirmed that DHHC6 can associate into higher order structures, possibly dimers but also higher order complexes, as opposed to DHHC 7 and 20 which appeared largely monomeric (Fig. 4D). The AAA mutant also formed higher order complexes but these were clearly different from those formed by the WT protein (Fig. 4D). Thus palmitoylation affects higher order assembly of DHHC6 and as well as localization of the protein in the ER.

### Palmitoylation controls DHHC6 activity

Finally we tested whether palmitoylation affects DHHC6 activity, which was monitored by measuring palmitoylation of one of its targets, namely calnexin. Calnexin palmitoylation was significantly reduced in DHHC16 KO cells, as monitored by Acyl-RAC (Fig. 4E) and enhanced upon DHHC16 over-expression (Fig. 4F) indicating that palmitoylation indeed modulates DHHC6 activity.

We next analysed the importance of the different palmitoylation sites. For this we overexpressed WT and DHHC6 mutants in Hela cells. Over-expression of WT DHHC6 led to 2.6 fold increase in calnexin palmitoylation, the background level being due to endogenous DHHC6 (Fig. 4GH). Overexpression of DHHC6 AAA had no effect, confirming the importance of palmitoylation for activity. The ACC, AAC and ACA showed very limited activity, as opposed to the CAA, CCA and CAC mutants (Fig. 4GH). Together these experiments indicate that palmitoylation of Cys-328 confers the highest activity to DHHC6.

### Model of the DHHC6 palmitoylation system

The results presented in the previous paragraphs show that DHHC6 can be palmitoylated on 3 sites, that the protein undergoes cycles of palmitoylation-depalmitoylation and that palmitoylation strongly influences assembly, localization, stability and function of the enzyme. The presence of 3 palmitoylation sites leads to the potential existence of 8 species: from fully unoccupied Cys-sites, noted C^000^, to full occupancy, noted C^111^, in the order of Cys-328, Cys-329 and Cys-343 in the exponent (Fig. 5A). To understand the dynamics of the inter-conversion between the 8 species, as well as derive hypothesis about the role of the three sites, we developed a mathematical model (Table S1). Modelling was performed as an open system, including protein synthesis, and degradation of all species (Fig. 5A and Table S2). Synthesis of DHHC6 first leads to an unfolded species (U). Folded DHHC6 protein is initially in the C^000^ form, which can undergo palmitoylation on any of the 3 sites leading to C^100^, C^010^, C^001^. These single palmitoylated species can undergo a second and a third palmitoylation event first leading to C^110^, C^101^, C^011^ and finally to C^111^. Each palmitoylation reaction is mediated by DHHC16 and each depalmitoylation catalysed by APT2. A competition term between the sites was implemented in the enzymatic kinetics, as previously for the modelling of palmitoylation of the ER chaperone calnexin (Dallavilla et al, 2016). The model includes first-order degradation rates for each species, with different rate constants. The rate expressions, the parameters and the assumptions used in the development of the model are described in detail in the Material and Methods and the Supplementary Information (Tables S1-3),

**Figure 5.**
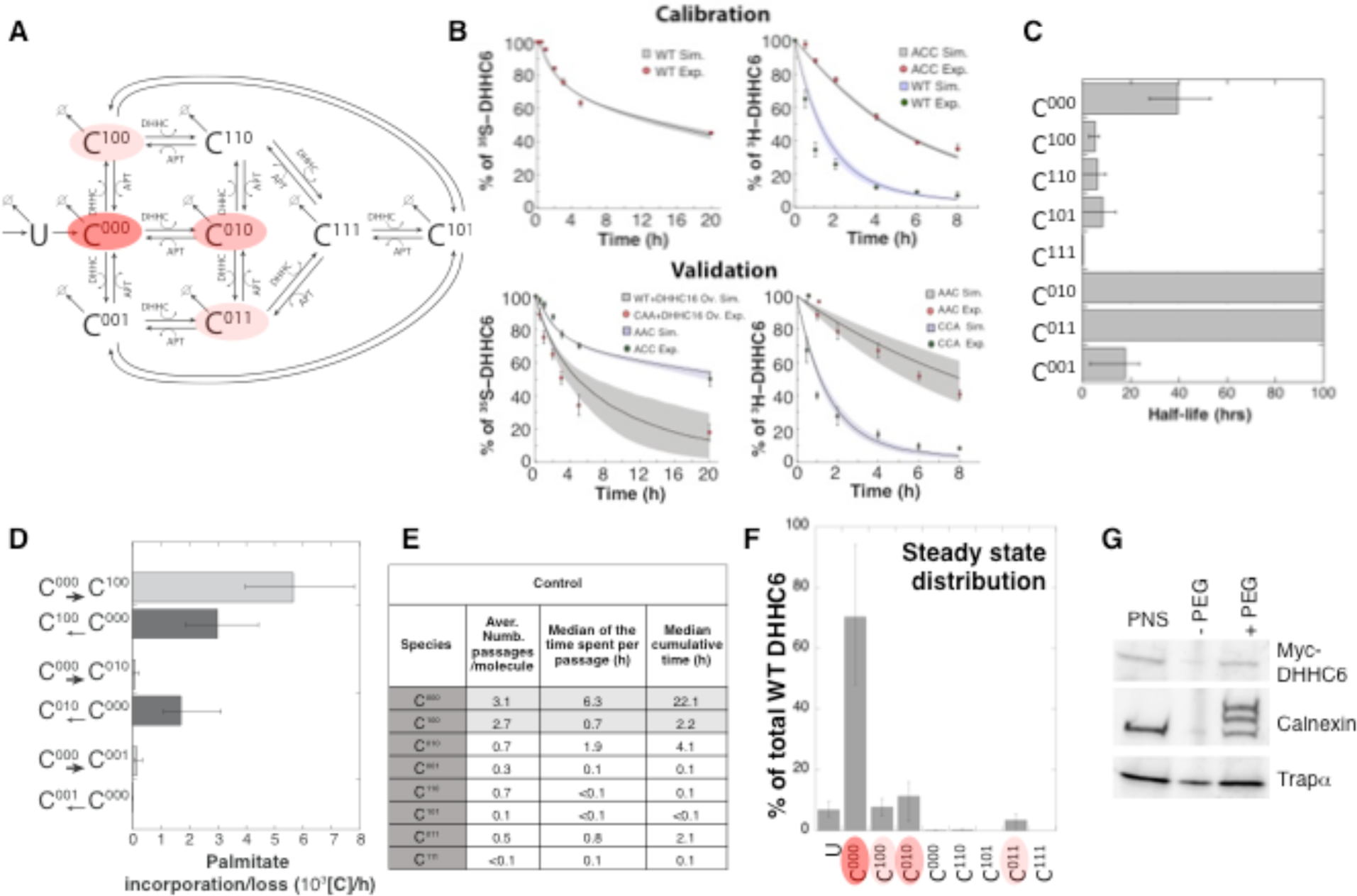
Modelling and analysis of DHHC6 palmitoylation. **A.** Network topology of the DHHC6 palmitoylation model. First we have a phase of synthesis of the unfolded peptide (U). The protein goes through a process of folding and membrane embedding, ending in the fully folded form (C^000^). The three sites can then be palmitoylated by DHHC16, the first palmitoylation can occur on each site, C^100^ C^010^ and C^001^ denote palmitoylation on the first, second or third site respectively. DHHC6 can then undergo another palmitoylation step acquiring two palmitates; C^110^, C^101^ and C^011^ denote the double palmitoylated enzyme. From each of the three double palmitoylated states, DHHC6 can be modified one last time becoming fully palmitoylated (C^111^). We consider that DHHC6 degradation can happen in any state. Highlighted in red are the most abundant species in steady state. **B.** Part of the calibration and validation sets used for parameter estimation. In the top graphs we show 3 curves that were used during parameter estimation with genetic algorithm to evaluate the goodness of fit of the parameters generated. Other curves used for the validation are shown in Fig. 5-figure supplement 1. The four curves in the bottom graphs are part of the data that were used for validating the predictions on experiments that were not used for parameter calibration. Circles represent experimental data while the solid lines are the output of the model after optimization. Since we have 152 different sets of optimal parameters, the shadows behind the lines represent the 1^st^ and 3^rd^ quartile of the 152 model outputs. All other curves used for parameter estimation can be found in Fig. 5-figure supplement 2. **C.** Half-life of the different DHHC6 palmitoylation states estimated from the decay rate constants of the model after optimization. The half-life was calculated as: ln(2)/kd_i_. Where kd_i_ is the decay rate constant of the i-th palmitoylation state. **D.** Palmitoylation and depalmitoylation fluxes for the six steps of single palmitoylation/depalmitoylation in steady state, which represent the fluxes of palmitate incorporation and loss on the first, second and third site during the first palmitoylation event. Fluxes in steady state for all reactions are shown in Fig. 5-Figure supplement 3. **E.** Single molecule tracking with stochastic simulations. The table shows the average number of passage per molecule for each state of the model, along with the median and the cumulative median of the time spent in each state. These data were obtained from the analysis of 10’000 stochastic simulations. **F.** Prediction of the steady state distribution of DHHC6 WT palmitoylation species. **G.** Stoichiometry of DHHC6 palmitoylation in Hela cells. Hela cells were transfected with plasmids encoding WT myc-DHHC6 constructs for 24h. **C.** Protein lysates were processed for the APEGS assay. PEG-5k was used to label transfected myc-DHHC6 and endogenous protein (PEG+), PEG-lanes indicate the negative controls. The samples were analysed by western blotting with anti-myc, anti-calnexin, anti-TRAP alpha antibodies.

With the aim of first calibrating and subsequently validating the model, we generated the following kinetic datasets. We performed: 1) metabolic ^35^SCys/Met pulse-chase experiments, with pulses of either 20 min or 2 hrs, to monitor the turnover of newly synthesized proteins, for WT and the different single, double and triple cysteine mutants; 2) ^3^H palmitate incorporation into WT and cysteine mutants; 3) palmitate loss, monitored by ^3^H palmitate pulse-chase experiments, for WT and mutants. Some of these experiments were in addition performed upon over expression of DHHC16 or siRNA of APT2.

We used the ^35^S-decay data for WT, ACC, CAA and AAA, the WT ^3^H-palmitate incorporation data and the ^3^H-palmitate release data for WT, ACC and CAA, to calibrate the model. During parameter estimation, these calibration datasets were fitted simultaneously, as multi-objective optimization problem with as many fitness functions as experiments. As often for optimization problems, the output of the algorithm is a local Pareto set of solutions. These are equally optimal with respect to the fitness functions in the sense that for each set of parameters, none of the objective functions can be improved in value without deteriorating the quality of the fitness for some of the other objective values. We therefore employed a stochastic optimization method to generate a population of models consistent with the calibration experiments. From a population of 10’000 models, we selected 152 based on the scores of the objective functions that fitted the experimental data most accurately (Fig. 5-figure supplement 1). The pool of selected models was subsequently used for the simulations and analyses. Importantly, all predictions were obtained by simulating each model independently. Outputs of all models were averaged and standard deviations with respect to the mean were calculated (Fig. 5B).

The remaining experiments, not used to calibrate the model, were used to validate it. Importantly, as can be seem in Fig. 5B and Fig. 5-figure supplement 2, the model reliably predicted the results of these experiments, indicating that it accurately captures the DHHC6 palmitoylation system.

### Dynamics and consequences of site-specific DHHC6 palmitoylation

The model was first used to estimate the half-lives of the different species. C^000^ is predicted to have a half-life of approx. 40 hrs (Fig. 5C and Table S4), consistent with the decay measured experimentally for the AAA triple mutant (Fig. 5-figure supplement 2). The presence of palmitate on Cys-328 is predicted to strongly accelerate protein turnover, irrespective of the occupancy of the other sites, with *t*_1/2_=5 hrs for C^100^ and *t*_1/2_=0.3 hrs C^111^ (Fig. 5C and Table S4). In contrast, palmitate on Cys-329 is predicted to have a stabilizing effect with *t*_1/2_> 100 hrs for C^010^. Finally, palmitate on Cys-343 would have a moderately destabilizing effect with *t*_1/2_= 18 hrs for C^001^ (Fig. 5C). Thus, consistent with the experimental observations, the model indicates that palmitoylation strongly affects DHHC6 turnover in a site dependent manner.

We next analysed the dynamics of the system. We first determined the palmitoylation and depalmitoylation fluxes. The major fluxes through the system are from C^000^ to C^100^, backwards from C^100^ to C^000^ and to a lesser extend from C^010^ to C^000^ (Fig. 5D). We then derived a stochastic formulation of the model, as we have previously done for calnexin palmitoylation (Dallavilla et al, 2016) (Suppl. Information, Tables S6-8). We performed 10’000 simulations to track single proteins in the system. DHHC6 molecules were predicted to spend most of their time, by far, in the C^000^ state (Fig. 5E). Consistent with the flux analysis, each DHHC6 molecule was on average 2.7 times in the C^100^ state (Fig. 5E), and remained in that state for about 0.7 hrs (median value). Seven out of 10 molecules also explored the C^010^ state for almost 2 hrs and more briefly the C^011^ state.

Estimation of the steady state distribution of species, under our experimental setting, indicated that the DHHC6 cellular population is about 70% unpalmitoylated and 20 % palmitoylated, the most abundant species being C^010^ and C^011^ (Fig. 5F). We tested this prediction experimentally by performing a PEGylation assay. This is a mass-tag labelling method that consists in replacing the palmitate moiety with PEG following disruption of the thioester bond with hydroxylamine, resulting in a mass change detectable by electrophoresis and Western blotting (Percher et al, 2016; Yokoi et al, 2016). As a control we analysed calnexin, which has 2 palmitoylation sites and migrated as expected as 3 bands, corresponding to the non-, single and dual palmitoylated forms (Fig. 5G).

For DHHC6, only the non-palmitoylated form was detected, consistent with the prediction (Fig. 5FG). Indeed given the dynamic range and sensitivity of Western blots, a band with a ≈7-fold lower intensity than the C^000^ DHHC6 band would not be detectable. The model altogether predicts that under our experimental conditions, each DHHC6 molecule undergoes multiple rounds of palmitoylation-depalmitolyation during its life cycle mostly on Cys-328. Only a subset of molecules acquire palmitate on the two other sites. At any given time, approximately 60% of the molecules are not acylated.

### Multi-site palmitoylation prevents large fluctuations in DHHC6 protein levels

Analysis of the activity of the DHHC6 mutants (Fig. 4H) showed that the most active variants are those with a cysteine at position 328, suggesting that palmitoylation at this position influences palmitoyltransferase activity. However palmitoylation at this site also targets DHHC6 to ERAD and thus leads to a decrease in the cellular content of DHHC6 (Fig. 3D). These effects raise the question of how cells can increase cellular DHHC6 activity.

We first predicted the consequences of DHHC6 hyperpalmitoylation. Calculations were performed under conditions of DHHC16 overexpression and of APT2 silencing, simulated by an absence of APT2. DHHC16 overexpression led to a major shift in species distribution, C^011^ then representing 60% of the population (Fig. 6A). Interestingly, the overall DHHC6 content dropped by only 15% (Table S5). APT2 silencing in contrast led to a 72% drop in DHHC6 content (Table S5). Since removing all depalmitoylating activity is likely to freeze the system, we also tested a decrease of APT2 to 10% of normal levels. The DHHC6 content rose by 20% and as upon DHHC16 overexpression C^011^ became the most abundant form (Fig. 6A). We made similar calculations for the CAA mutant. Under control conditions, the total CAA content is predicted to be 84% of WT, and mostly in the non-palmitoylated state (Fig. 6B). The CAA cellular content is however predicted to drop to 32% of WT control levels upon DHHC16 overexpression (Fig. 6B). All together, these predictions suggest that the presence of 3 palmitoylation sites, as opposed to just Cys-328, renders the DHHC6 protein content more robust to changes in DHHC16 activity.

**Figure 6.**
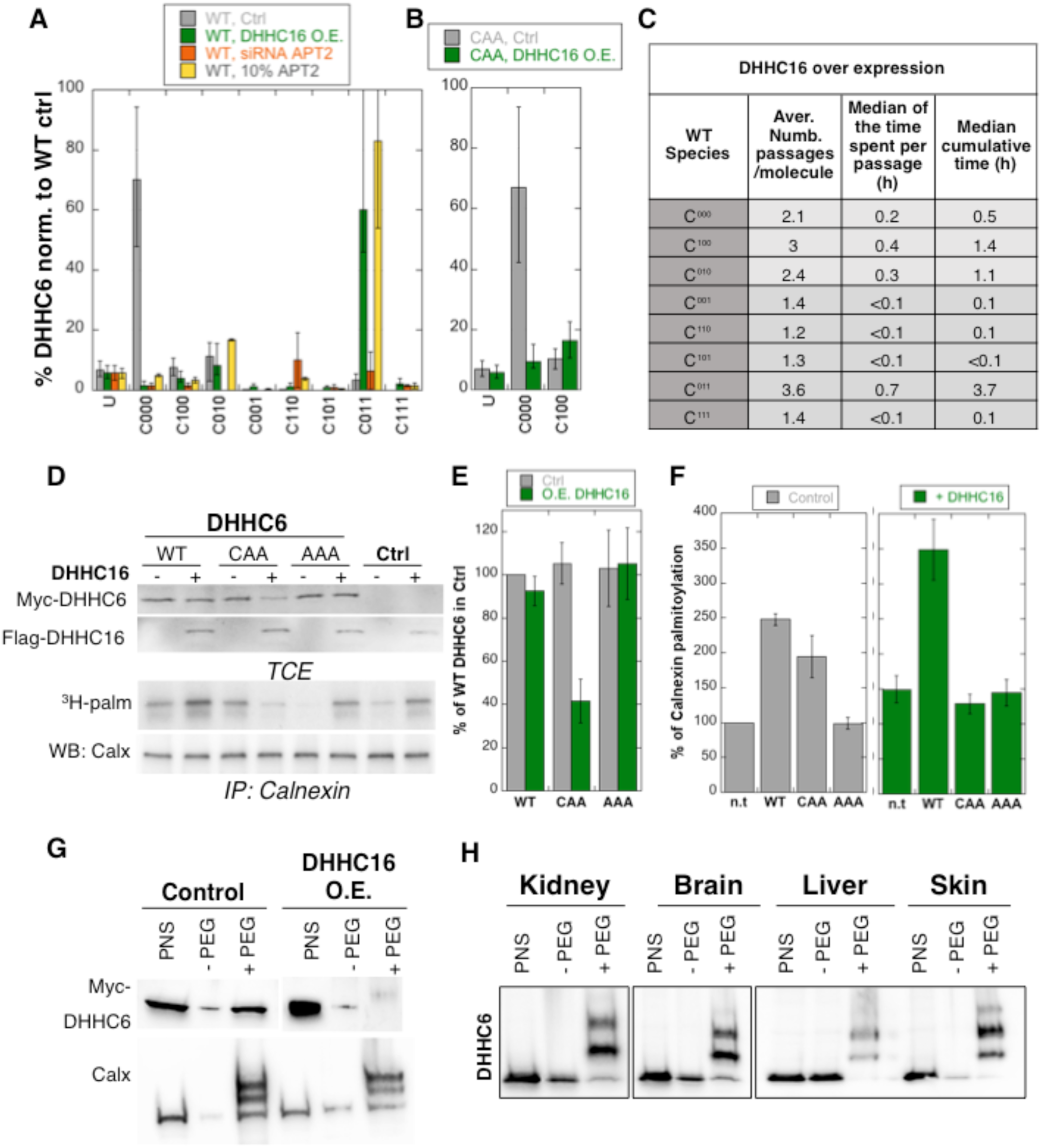
Importance of multiple palmitoylation sites. **A.** Single molecule tracking with stochastic simulations upon DHHC16 overexpression. The table shows the average number of passage per molecule in each state of the model, along with the median and the cumulative median of the time spent in each state. As for control experiments this numbers were obtained by averaging the results of 10’000 independent stochastic simulations. **B.** Steady state DHHC6 WT species distribution under different conditions (control, DHHC16 Overexpression, APT2 silencing and APT2 at 10% level of expression with respect to WT condition). All the data are scaled with respect to the total abundance of DHHC6 WT under control condition. **C.** Steady state distribution of the DHHC6 CAA mutant under control condition or DHHC16 overexpression. All the data are scaled with respect to the total abundance of DHHC6 WT in normal condition. **D.** Calnexin palmitoylation upon DHHC6 and DHHC16 overexpressed. Hela cells were transfected with control plasmid (Ctrl) or plasmids encoding WT or mutants myc-DHHC6 in the presence or not of FLAG-DHHC16 for 24h. Cells were then metabolically labelled 2 hours at 37°C with ^3^H-palmitic acid. Proteins were extracted, TCE isolated, and immunoprecipitated with anti-calnexin antibodies, subjected to SDS-PAGE and analysed by autoradiography (^3^H-palm), quantified using the Typhoon Imager or by immunoblotting with anti-calnexin, anti-myc or anti-flag antibodies. **E.** Quantification of DHHC6 levels in total cell extracts with or without DHHC16 overexpression. The calculated value for WT DHHC6 under control condition was set to 100%. All DHHC6 mutants with or without DHHC16 were expressed relative to this (n=4, errors represent standard deviations). **FG.** Quantification of ^3^H-palmitic acid incorporation into endogenous calnexin with DHHC6 in the absence (**F**) or presence of DHHC16 (**G**). The calculated value of ^3^H-palmitic acid incorporation into calnexin was set to 100% for control plasmid (Ctrl) and all DHHC6 mutants were expressed relative to this. N=4. **H.** Stoichiometry of DHHC6 palmitoylation in Hela cells with DHHC16 overexpressed. Hela cells were transfected with plasmids encoding WT myc-DHHC6 with Flag-DHHC16 constructs for 24h. Protein lysates were processed for the APEGS assay. PEG-5k were used for labelling of transfected myc-DHHC6 (PEG+), PEG-lanes indicate the negative controls. The samples were analysed by western blotting with anti-myc. **I.** Adult mouse tissues were processed for the APEGS assay. PEG-5k were used for labelling of endogenous DHHC6 (PEG+), PEG-lanes indicate the negative controls. The samples were analysed by western blotting with rabbit anti-DHHC6.

Stochastic simulations predicts that overexpression of DHHC16 drastically increased the dynamics of the network (Fig. 6C vs. 5E). Indeed, all DHHC6 molecules explored all the palmitoylation states, with extremely short residence times in each (Fig. 6C). Interestingly, flux analysis indicated that the abundance of C^011^ was primarily due to two 4-step paths: both started with C^000^ to C^100^, followed by palmitoylation on either of the two remaining sites, subsequent depalmitoylation of Cys-328, and finally palmitoylation of the remaining site (Fig. 5-figure supplement 3). When analysing 10’000 simulations, 22’000 events of palmitoylation-depalmitoylation emanated from C^011^ whereas only 4’000 events occurred between C^000^ and C^100^. Thus when activity of DHHC16 is high, C^011^ appears to be the hub of the system. C^011^ is also which is the most stable state for DHHC6, explaining that the protein levels are stable.

We next sought to validate these predictions experimentally. Western blot analysis of protein abundance showed that WT DHHC6 protein levels drastically dropped upon siRNA of APT2 (Fig. 4B), but not upon DHHC16 overexpression (Fig 6DE). DHHC16 overexpression however led to a 60% drop in CAA expression. As expected, expression of AAA was not affected (Fig. 6DE).

### Multi-site palmitoylation regulates DHHC6 activity

We next analysed the importance of multiple palmitoylation sites on DHHC6 activity, again monitored by measuring calnexin palmitoylation. DHHC16 overexpression enhanced the ability of WT DHHC6 to palmitoylate calnexin, whereas it essentially abolished the activity of CAA (Fig. 6 DFG). Note that the “background” calnexin palmitoylation level was increased due to the activation of endogenous DHHC6 by DHHC16 over expression. Using PEGylation, we confirmed that DHHC16 overexpression leads to an increase of the palmitoylated DHHC6 species (Fig. 6H).

The above experimental analyses show that in Hela cells under control tissue culture conditions, DHHC6 is mainly in the non-palmitoylated form. When DHHC6 PEGylation experiments were performed on different mouse tissues, we however found that DHHC6 is far more palmitoyled, suggesting that palmitoylation-mediated DHHC6 activity is higher *in vivo*.

## MATERIAL AND METHODS

### Cell lines

Hela cells (ATCC) were grown in complete modified Eagle’s medium (MEM, Sigma) at 37°C supplemented with 10% fetal bovine serum (FBS), 2mM L-Glutamine, penicillin and streptomycin. For the DHHC6 knockdown cell lines, Hela cells were transfected with shRNA against DHHC6 gene (target sequence in 3’UTR: 5'-CCTAGTGCCATGATTTAAA-3’) or with shRNA control against firefly luciferase gene (target sequence: 5’-CGTACGCGGAATACTTCGA-3’). The transfected cells were selected by treatment with 3µg/ml puromycin. HAP1 Wild type WT and knockout cell lines were purchased from Horizon Genomics (Vienna, Austria). The DHHC6 clone (13474-01) contains a 5bp deletion in exon 2 (NM_022494) and the DHHC16 clone (36523-06) contains a 2bp insertion in exon 2 (NM_198043). HAP1 cells were grown in complete Dulbeccos MEM (DMEM, Sigma) at 37°C supplemented with 10% fetal bovine serum (FBS), 2mM L-Glutamine, penicillin and streptomycin.

### Antibodies and Reagents

The following primary antibodies are used: Mouse anti-Actin (Millipore, MAB 1510), Mouse anti-Myc 9E10 (Covance, MMs-150R), Mouse anti-Ubiquitin (Santa Cruz, sc-8017), Mouse anti-GFP (Roche, 11814460001), Rabbit anti-DHHC6 (Sigma, SAB1304457), Mouse anti-Transferrin Receptor (Thermo Scientific, 136800), Mouse anti-Flag M2 (Sigma, F3165), Rabbit anti-ANTXR1 (Sigma, SAB2501028), Rabbit anti-TRAPalpha (Abcam, ab133238), Rabbit anti-Flotillin1 were produced in our laboratory, Mouse anti-CLIMP63 (Enzo, ALX-804-604), Mouse anti-Calnexin (MAB3126), Rabbit anti-GP78 AMFR(Abnova, PAB1684), Rabbit anti-IP3R (Cell signaling, 85685). The following beads were used for immunoprecipitation: Protein G Sepharose 4 Fast flow (GE Healthcare, 17-0618-01), anti-Myc affinity gel (Thermo Scientific, 20169), anti-Flag affinity gel EZview M2 (Sigma, F2426). Drugs were used as follows: Bafilomycin A1 at 100 nM (Sigma, B1793), MG132 at 10 µM (Sigma, C2211), Hydroxylamine at 0.5 M (Sigma, 55459), mPEG-5k at 20mM (Sigma, 63187), N-ethylmaleimide NEM at 20mM (Thermo Scientific, 23030), Tris-2- carboxyethyl-phosphine hydrochloride TCEP at 10 mM (Thermo Scientific, 23225), Methyl methanethiosulfonate MMTS at 1.5% (Sigma, 208795), Puromycin at 3µg/ml (Sigma, P9620).

### Transfection and siRNA experiments

Human Myc -DHHC6, Myc-DHHC6-C328A (ACC), Myc-DHHC6-C329A (CAC), Myc-DHHC6-C343A (CCA), Myc-DHHC6-C328A-C329A (AAC), Myc-DHHC6-C326A-C343A (ACA), Myc-DHHC6-C329A-C343A (CAA), Myc-DHHC6-C328A-C329A-C343A, Myc-DHHC6-R361Q, Myc-DHHC6-R361A, Myc-DHHC6-Del-K410-K411-N412-R413, Myc-DHHC6-Del-D126-H127-H128-C129 were cloned in pcDNA3. Human GFP-DHHC6 and all cyteine mutants mentioned above were also cloned in peGFP. Human FLAG-DHHC16 was cloned in pCE-puro-3xFLAG, human Myc-DHHC16 was cloned in pCE puro-his-myc. All other human Myc-DHHCs were cloned in pcDNA3.1 (provided by the Fukata lab). mCitrine fusions of APTs were inserted into pcDNA3.1-N1 (provided by Bastiaens lab, (Vartak et al, 2014)). mCitrine APT1-S119A, mCitrine APT2-S121A, mCitrine-APT1-C2S, mCitrine-APT2-C2S were cloned in pcDNA3.1-N1. For control transfection, we used an empty pcDNA3 plasmid. Plasmids were transfected into HeLa cells for 24h (2µg cDNA/9.6cm^2^) plate using Fugene (Promega).

For gene silencing, Hela cells were transfected for 72 h with 100pmol/9.2cm2 dish of siRNA using interferin (Polyplus) transfection reagent. As control siRNA we used the following target sequence of the viral glycoprotein VSV-G: ATTGAACAAACGAAACAAGGA. siRNA against human genes were purchased from Qiagen (DHHC6 target sequences: 1-GAGGTTTACGATACTGGTTAT, 2-TAGAAGGTGTTTCAAGAATAA, DHHC16 target sequences: 1-CTCGGGTGCTCTTACCTTCTA, 2-TAGCATCGAAAGGCACATCAA; APT1 target sequence: AACAAACTTATGGGTAATAAA; APT2 target sequence: AAGCTGCTGCCTCCTGTCTAA, all other DHHC target sequences were previously tested (Lakkaraju et al, 2012)).

### Real-time PCR

For Hela cells, RNA was extracted from a six-well dish using the RNeasy kit (Qiagen). 1 mg of the total RNA extracted was used for the reverse transcription using random hexamers and superscript II (Thermo Scientific). A 1:40 dilution of the cDNA was used to perform the real-time PCR using SyBr green reagent (Roche). mRNA levels were normalized using three housekeeping genes: TATA-binding protein, β-microglobulin and β-glucoronidase. Total RNA of different mouse tissues were extracted using the RNeasy kit (Qiagen) after solubilization with TissueLyser II (Qiagen).

### Radiolabelling experiments

For the ^35^S-metabolic labeling, the cells were starved in DMEM HG devoid of Cys/Met for 30 minutes at 37°C, pulsed with the same medium supplemented with 140 µCi of ^35^S Cys/Met (American Radiolabeled Chemicals, Inc.) for the indicated time, washed and incubated in DMEM complete medium for the indicated time of chase (Abrami et al, 2008) before immunoprecipitation. To detect palmitoylation, Hela cells were transfected or not with different constructs, incubated for 2 hours in IM (Glasgow minimal essential medium buffered with 10mM Hepes, pH 7.4) with 200 µCi/ml 3H palmitic acid (9,10-^3^H(N)) (American Radiolabeled Chemicals, Inc.). The cells were washed, incubated in DMEM complete medium for the indicated time of chase, or directly lysed for immunoprecipitation with the indicated antibodies.

For all radiolabelling experiments, after immunoprecipitation, washes beads were incubated for 5 min at 90°C in reducing sample buffer prior to 4-12% gradient SDS-PAGE. After SDS-PAGE, the gel are incubated in a fixative solution (25% isopropanol, 65% H2O, 10% acetic acid), followed by a 30 min incubation with signal enhancer Amplify NAMP100 (GE Healthcare). The radiolabeled products were revealed using Typhoon phosphoimager and quantified using the Typhoon Imager (ImageQuanTool, GE Healthcare).

### Immunoprecipitation

For immunoprecipitation, cells were washed 3 times PBS, lysed 30min at 4°C in the following Buffer (0.5% Nonidet P-40, 500 mM Tris pH 7.4, 20 mM EDTA, 10 mM NaF, 2 mM benzamidin and protease inhibitor cocktail (Roche)), and centrifuged 3min at 5000 rpm. Supernatants were subjected to preclearing with G sepharose beads prior immunoprecipitation reaction. Supernatants were incubated overnight with the appropriate antibodies and G Sepharose beads.

### Post nuclear supernatants and ACYL-RAC

HeLa cells were harvested, washed with PBS, and homogenized by passage through a 22G injection needle in HB (HB: 2.9 mM imidazole and 250 mM sucrose, pH 7.4) containing a mini tablet protease inhibitor mixture (Roche). After centrifugation, the supernatant was collected as PNS (Post Nuclear Supernatant).

Protein S-palmitoylation was assessed by the Acyl-RAC assay as described (Werno & Chamberlain, 2015), with some modifications. A fraction of the PNS was saved as the input. Hela PNS were lysed in buffer (0.5% Triton-X100, 25 mM HEPES, 25 mM NaCl, 1 mM EDTA, pH 7.4 and protease inhibitor cocktail). In order to block free SH groups with S-methyl methanethiosulfonate (MMTS), 200µl of blocking buffer (100 mM HEPES, 1 mM EDTA, 87.5 mM SDS and 1.5% (v/v) MMTS) was added to cell lysate and incubated for 4 h at 40°C. Subsequently, 3 volumes of ice-cold 100% acetone was added to the blocking protein mixture and incubated for 20 minutes at -20 °C and then centrifuged at 5,000 × g for 10 minutes at 4 °C to pellet precipitated proteins. The pellet was washed five times in 1 ml of 70% (v/v) acetone and resuspended in buffer (100 mM HEPES, 1 mM EDTA, 35 mM SDS). For treatment with hydroxylamine (HA) and capture by Thiopropyl Sepharose® beads, 2 M hydroxylamine was added together with the beads (previously activated for 15 min with water) to a final concentration of 0.5 M hydroxylamine and 10% (w/v) beads. As a negative control, 2 M Tris was used instead of hydroxylamine. These samples were then incubated overnight at room temperature on a rotate wheel. The beads were washed, the proteins were eluted from the beads by incubations in 40 µl SDS sample buffer with beta-mercapto-ethanol for 5 minutes at 95 °C. Finally, samples were submitted to SDS-PAGE and analyzed by immunoblotting.

### APEGS assay

The level of protein S-palmitoylation was assessed as described (Yokoi et al, 2016), with minor modifications. Hela cells were lysed with the following buffer (4% SDS, 5 mM EDTA, in PBS with complete inhibitor (Roche)). After centrifugation at 100,000 × *g* for 15 min, supernatant proteins were reduced with 25 mM TCEP for 1 h at 55°C or at room temperature (RT), and free cysteine residues were alkylated with 20 mM NEM for 3 h at RT to be blocked. After chloroform/methanol precipitation, resuspended proteins in PBS with 4% SDS and 5 mM EDTA were incubated in buffer (1% SDS, 5 mM EDTA, 1 M NH_2_OH, pH 7.0) for 1 h at 37°C to cleave palmitoylation thioester bonds. As a negative control, 1 M Tris-HCl, pH7.0, was used. After precipitation, resuspended proteins in PBS with 4% SDS were PEGylated with 20 mM mPEGs for 1 h at RT to label newly exposed cysteinyl thiols. As a negative control, 20 mM NEM was used instead of mPEG (-PEG). After precipitation, proteins were resuspended with SDS-sample buffer and boiled at 95°C for 5 min. Protein concentration was measured by BCA protein assay.

### BLUE Native PAGE

The PNS of Hela cells were extracted. The proteins were lysed in 1% Digitonin and passed through a 26g needle, incubated on ice 30 minutes, spin 16 000g for 30 minutes and run following the manufacturer instructions on the Novex Native PAGE Bis-tris gel system (ThermoFisher).

### Immunofluorescence staining and fluorescence microscope

HeLa cells were seeded on 12mm glass coverslips (Marienfeld GmbH, Germany) 24hr prior to transfection. PAT6-myc plasmids were transfected using Fugene (Promega,USA) for 48hrs. Cells were then fixed using 3% paraformaldehyde for 20min at 37°C, quenched 10min with 50mM NH_4_Cl at RT and permeabilized with 0.1% TX100 for 5min at RT and finally blocked overnight with 0.5% BSA in PBS. Cells were washed 3x with PBS in between all the steps. Cells were then stained with anti-myc and anti-BiP antibodies for 30min at RT, washed and incubated again 30min with their corresponding fluorescent secondary antibodies. Finally cells were mounted on glass slide using mowiol. Imaging was performed using a confocal microscope (LSM710, Zeiss, Germany) with a 63x oil immersion objective (NA 1.4).

### Core model of DHHC6 palmitoylation

The DHHC6 palmitoylation model (Fig. 5A) was designed following the approach elaborated previously for calnexin (Dallavilla et al, 2016). The core model is based a previously described protein phosphorylation model used in (Goldbeter & Koshland, 1981). The set of reaction described by Goldbeter was used to model a single palmitoylation event. Multiple palmitoylation events were modelled replicating this subunit for each reaction.

In the Goldbeter study, the model was mathematically described using mass action terms. Here, due to the presence of multiple modification events, which require the definition of a consistent number of parameters, we described model reactions using so-called “total quasi-steady state approximation” (tQSSA) (Pedersen et al, 2008). With respect to the standard quasi-steady state approximation (QSSA), tQSSA is valid also when the enzyme-substrate concentrations are comparable (Borghans et al, 1996). DHHC6 is palmitoylated by DHHC16. Due to this fact, the use of tQSSA is justified since different proteome studies suggest that DHHC enzymes have similar concentrations (Beck et al, 2011; Merrick et al, 2011; Nagaraj et al, 2011; Yang et al, 2010; Yount et al, 2010). The step-by-step application of the tQSSA approximation to a palmitoylation model is described in (Dallavilla et al, 2016).

The model can be divided in two parts; we first have a phase of synthesis of DHHC6, which initially is unfolded (U in Fig. 5A) and subsequently reaches the folded initially non-palmitoylated (C^000^) form. Each site can subsequently be palmitoylated leading to C^100^ C^010^ and C^001^. These species can then undergo a second palmitoylation step leading to C^110^, C^101^ and C^011^, and a third leading to C^111^. Since palmitoylation is reversible, palmitate can be removed from each of the sites, from all of the species.

DHHC6 model is based on the following assumption:

- DHHC6 can be degraded in each of its states.
- DHHC6 is present in similar concentrations with respect to the modifying enzyme DHHC16 (Beck et al, 2011; Merrick et al, 2011; Nagaraj et al, 2011; Yang et al, 2010; Yount et al, 2010).
- Acyl protein thioesterases (APTs) are more abundant than DHHC6 (Beck et al, 2011; Merrick et al, 2011; Nagaraj et al, 2011; Yang et al, 2010; Yount et al, 2010)
- The three palmitoylation sites may have different affinities with respect to the palmitoylation/depalmitoylation enzymes, therefore we adopted separated Kms and V_max_s for the different sites.
- All the palmitoylation steps are reversible. APT catalyses the depalmitoylation steps.
- Palmitate was considered to be available in excess.

A detailed account of all reactions and differential equations in the model is given in Tables S1-3.

### Parameterization of the model

Since no kinetic data on DHHC6 palmitoylation is available, all the parameters of the model were estimated using a genetic algorithm. For the parameter optimization, multiple datasets coming from experimental results were considered. Time course labelling experiments were performed in order to characterize the dynamics of DHHC6 synthesis/degradation and incorporation/loss of palmitate.

The types of data used, the description of the Genetic Algorithm (GA) and the step-by-step application of the optimization algorithm used to find values for the parameters are described in detail in (Dallavilla et al, 2016). In this paper we use the exact same procedure.

For the parameter estimation we used a calibration set that correspond to the data visible in (Fig. 5B and Fig.S5). The remaining part of the data (Fig.5B and S6) was used to validate the output of the model and to verify the accuracy of its prediction capabilities. Because of the presence of multiple objectives there does not exist a single solution that simultaneously optimizes each objective, so the algorithm provides as output a local Pareto set of solutions, which are equally optimal with respect to the fitness function we defined. From the Pareto set provided by the GA and in order to be more accurate and to reduce the variability in the output of the model, we selected the set of parameters that best fitted the calibration data. The procedure for the selection of the subset is described in (Dallavilla et al, 2016). In total, 152 different sets of parameter were selected at the end of the optimization.

The results of the optimization can be found in Table S3.

### Simulating the labeling experiments

Since the acuracy of each parameter set is evaluated by computing the distance between the output of the model and different experiments of the calibration dataset, we made use of a previously a established method to reproduce the different type of experiment in-silico (Dallavilla et al, 2016).

### Stochastic simulations

In order to measure the average palmitoylation time of DHHC6, we made use of a previously established method to perform *in-silico* single molecule tracking (Dallavilla et al, 2016).

### Conversion of deterministic parameters to stochastic

Parameter conversion from deterministic to stochastic model is needed to perform stochastic simulations (Dallavilla et al, 2016). This transformation involves a change of units, from concentration to number of molecules. A single assumption was added to those previously defined (Dallavilla et al, 2016), namely that the number of DHHC6 molecules per HeLa cell is in the order of 1600 molecules (Beck et al, 2011; Merrick et al, 2011; Nagaraj et al, 2011; Yang et al, 2010; Yount et al, 2010) Table S6 shows the parameters obtained through the conversion. The model design is shown through the stoichiometry matrix and the propensity function in Tables S7-S8.

## DISCUSSION

S-Palmitoylation is a reversible lipid modification that can control protein function in time and in space. The enzymes involved must therefore also be regulated. The DHHC family of palmitoyltransferases was identified almost 15 years ago, yet little is known about the mechanisms that control their activity. DHHC9 was found to require a co-factor, GCP16, for activity (Swarthout et al, 2005), as also found in yeast (Lobo et al, 2002). Similarly, DHHC6 was proposed to require Selenoprotein K to function (Fredericks & Hoffmann, 2015). Here we show that DHHC6 activity is controlled by palmitoylation in its C-terminal SH3_2-domain. In the non-palmitoylated form, DHHC6 has no detectable activity. Only upon modification by an upstream palmitoylatransferase, which we identified as DHHC16, does it acquire significant transferase activity. DHHC16, also known as Ablphilin 2 (Aph2), is, as DHHC6, expressed in many human tissues in particular in the heart, pancreas, liver, skeletal muscle (Zhang et al, 2006). Its name Ablphilin stems from its ability to interact with the non-receptor tyrosine kinase c-Abl at the ER surface (Li et al, 2002). DHHC16/Aph2 is essential for embryonic and postnatal survival, eye and heart development in mice (Zhou et al, 2015), for the proliferation of neural stem cells, where it in involved in the activation of the FGF/erk pathways (Shi et al, 2015) and was reported to play a role in DNA damage response (Cao et al, 2016). Our findings raise the possibility that some of these effects may involve DHHC6.

We found that DHHC6 palmitoylation is highly dynamic, involving Acyl Protein Thioesterase 2. APT2, which was recently found to be involved in cell polarity-mediated tumor suppression (Hernandez et al, 2017) but is otherwise poorly characterized, also undergoes palmitoylation by a yet to be determined palmitoyltransferase (Kong et al, 2013; Vartak et al, 2014).

DHHC6 palmitoylation can occur on three sites in its SH3_2 domain. We combine mathematical modelling, with mutagenesis of palmitoylation sites, expression levels of the involved enzymes (DHHC16 and APT2) and experimental determination of palmitoylation, depalmitoylation, metabolic pulse-chase experiments and activity determination, to understand the importance of the different DHHC6 palmitoylation species and the dynamics of their interconversion. In addition to the predictive power of mathematical modelling of biological systems, this study provides single molecule understanding, as opposed to population analysis.

We found that in tissue culture cells, under standard conditions, DHHC6 molecules spend more than 70% of their lifetime in the non-palmitoylated inactive C^000^ state (Fig. 5E). Each DHHC6 molecule does however undergo several rounds of palmitoylation-depalmitoylation on Cys-328 (Fig. 5E), leading to C^100^, the most active state of the enzyme. According to large-scale quantitative proteomics analyses, DHHC enzymes are typically present in less then 1000 copies per cell (Beck et al, 2011; Foster et al, 2003; Nagaraj et al, 2011). Together this information indicates that, at any given time, tissue culture cells contain a very low number of active DHHC6 enzymes. In these cells, target proteins do however undergo palmitoylation, and some of these targets are highly abundant proteins such as calnexin (half a million to a million copies per cell (Beck et al, 2011; Foster et al, 2003; Nagaraj et al, 2011)). Thus when active, DHHC6 appears be a very potent enzyme and our observations indicates that cells tend to avoid having too much of it. Indeed the C^100^ species of DHHC6 is either rapidly depalmitoylated or targeted to degradation via the ERAD pathway.

PEGylation analysis of mouse tissues indicate that *in vivo* DHHC6 palmitoylation is far more pronounced than in tissue cultured cells (Fig. 6H). This situation was mimicked experimentally by overexpression of DHHC16 and revealed the importance of having 3 palmitoylation sites, rather than just Cys-328. We indeed found that in the presence of 3 sites, the cellular DHHC6 activity could be enhanced by DHHC16 overexpression. Increase DHHC16 activity led to a shift of species distribution C^011^, the most stable form, becoming the most abundant. Altogether this study indicates that DHHC6 can adopt in 3 types of states: 1) stable and inactive (C^000^); 2) highly active, short-lived and rapidly turned over (C^100^); 3) moderately active, long lived and highly stable (C^010^ and C^011^). Such a regulatory system allows the cell to tightly control the activity of DHHC6. Why excessive DHHC6 activity is detrimental to a cell or an organism will require further studies.

By combining experimentation and the predictive power of data-driven mathematical modelling, we were able to obtain unprecedented insight into the dynamics of palmitoylation and the role and properties of single acylated species. We indeed found that depending on the site of modification both the activity and turnover of DHHC6 were strongly affected, indicating that palmitoylation has a more subtle and precise effect that merely increasing hydrophobicity, consistent with recent findings on single acylated species of Ras (Pedro et al, 2017). We moreover revealed that palmitoylation can occur in cascades, DHHC16 controlling the activity of DHHC6, which itself tunes the activity of key proteins of the ER and other proteins of the endomembrane system. Finally this study highlights the importance of acyl protein thioesterases in regulating the interconversion between palmitoylted species.

## AUTHOR CONTRIBUTIONS

Conceptualization, L.A., T. D., P.S., G.V.D.G., V.H.; Investigation, L.A., T. D., P.S., M.D., N.G., G.K.; Funding Acquisition, G.V.D.G., V.H.; Writing–Original Draft, L.A., T. D., P.S., G.V.D.G., V.H.; Writing–Review & Editing, G.S., M.D., N.G., G.K.; Supervision, G.S.; Resources, L.A., T. D., P.S., B.K.

## ACKNOWLEDGEMENTS

We thank L. Chamberlain and B. Martin for sharing their Acyl-Rac protocol, Y. and M. Fukata for the DHHC plasmids and the PEGylation protocol, P. Bastiaens for the APT plasmids and Sylvia Ho for generating the shRNAi DHHC6 construct.

The research leading to these results has received funding from the European Research Council under the European Union’s Seventh Framework Programme (FP/2007-2013) / ERC Grant Agreement n. 340260 - PalmERa’. This work was also supported by grants from the Swiss National Science Foundation (to G.v.d.G and to V.H.), and the Swiss SystemsX.ch initiative evaluated by the Swiss National Science Foundation (LipidX) (to G.v.d.G and to V.H.). T.D. is a recipient of an iPhD fellowship from the Swiss SystemsX.ch initiative.

## Supplementary Figure Legends

**Figure 1-figure supplement 1.**
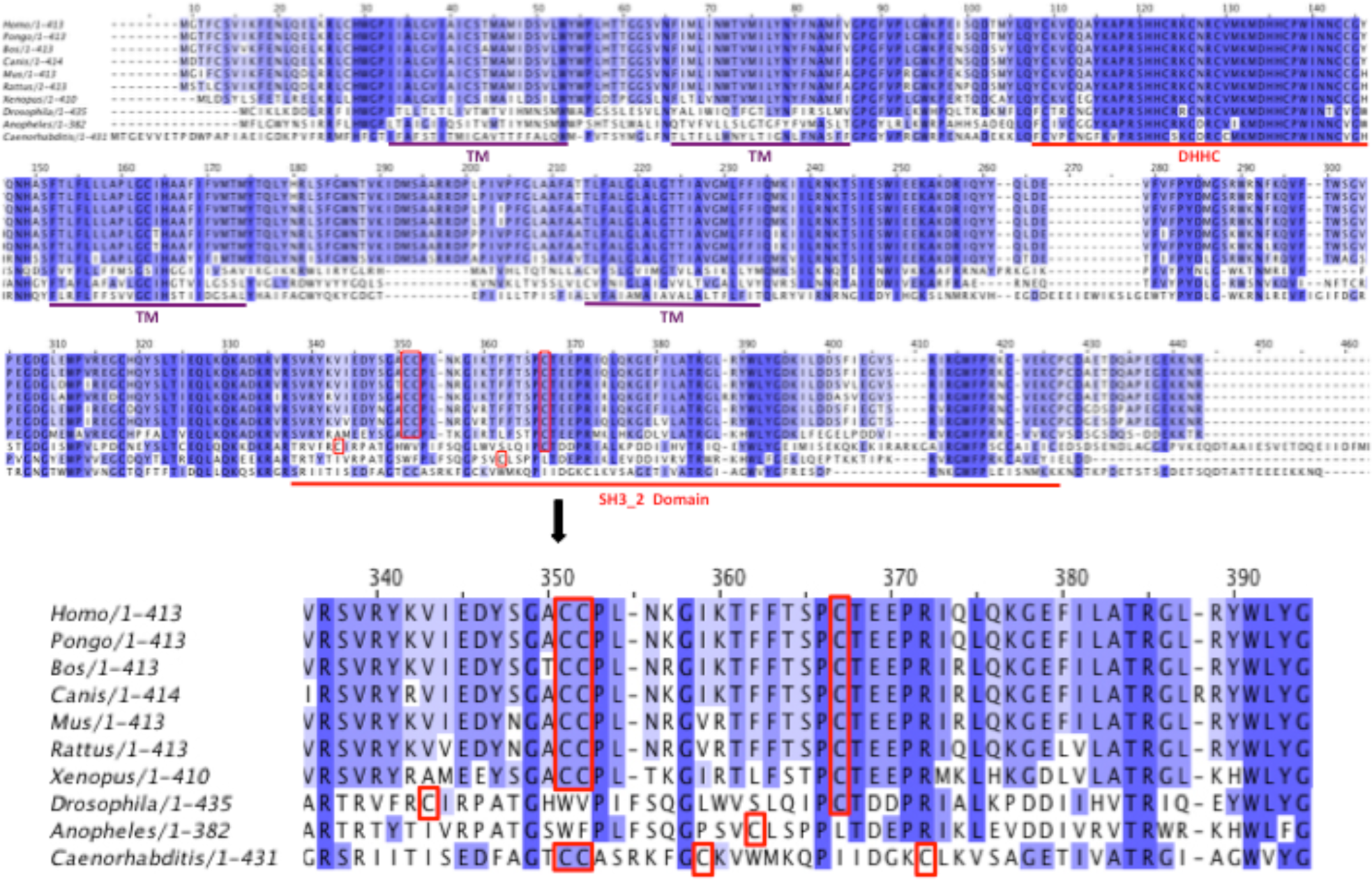
Alignment of DHHC6 sequences from different species. Identical amino acids are highlighted by dark blue. Palmitoylated cysteines are highlighted in red boxes. R361 is highlighted in green box. SH3-2 like domain and DHHC domains are underlined in red.

**Figure 2-figure supplement 1.**
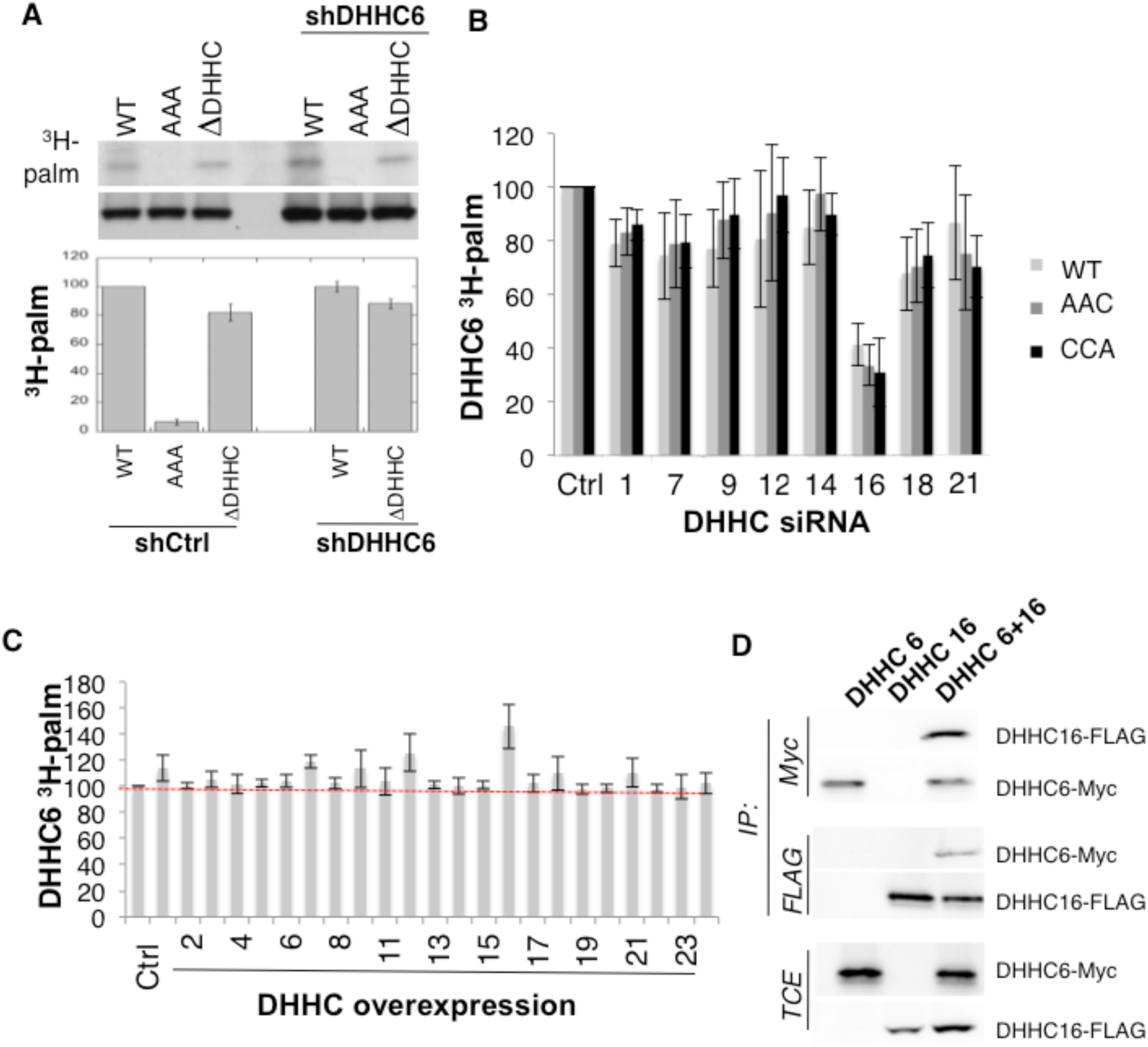
**A.** DHHC6 is not autopalmitoylated. Hela cells silenced with control lentiviruses or with shDHHC6 lentiviruses were transfected with plasmids encoding WT myc-DHHC6 or mutant myc-DHHC6. Cells were then metabolically labeled 2 hours at 37°C with 3H-palmitic acid. Proteins were extracted and immunoprecipitated with anti-myc antibodies, subjected to SDS-PAGE and analyzed by autoradiography (^3^H-palm), quantified using the Typhoon Imager or by immunoblotting with anti-myc antibodies. 3H-palmitic acid incorporation into DHHC6 values were calculated and were set to 100% for WT DHHC6 constructs and all DHHC6 mutants were expressed relative to this. N=4. **B.** All DHHC6 cysteine are palmitoylated by DHHC16. Hela cells were transfected with siRNA silencing indicated DHHC for 72h and with WT and cysteine mutants myc-tagged DHHC6 construct for the last 24h. Cells were then metabolically labeled 2 hours at 37°C with 3H-palmitic acid. Proteins were extracted, immunoprecipitated with myc antibodies and subjected to SDS-PAGE and analyzed by autoradiography, quantified using the Typhoon Imager or by immunoblotting with myc antibodies. ^3^H-palmitic acid incorporation into WT and mutants DHHC6 constructs were quantified and normalized to protein expression level. The calculated value of ^3^H-palmitic acid incorporation into WT or mutants DHHC6 was set to 100% for a non relevant siRNA (Ctrl) and all siRNA were expressed relative to this. N=3. **C.** Palmitoylation of WT DHHC6 after human DHHC overexpression. Hela cells were transfected with indicated DHHC constructs for 24h with WT myc-tagged DHHC6 construct. Cells were then metabolically labeled 2 hours at 37°C with 3H-palmitic acid. Proteins were extracted, immunoprecipitated with myc antibodies and subjected to SDS-PAGE and analyzed by autoradiography, quantified using the Typhoon Imager or by immunoblotting with myc antibodies. ^3^H-palmitic acid incorporation into WT DHHC6 constructs were quantified and normalized to protein expression level. The calculated value of 3H-palmitic acid incorporation into WT DHHC6 was set to 100% for a non relevant plasmid (Ctrl) and all DHHC were expressed relative to this. N=5. **D.** Co-immunoprecipitation of DHHC6 with DHHC16. Hela cells were transfected with plasmids encoding WT myc-DHHC6 and flag-tagged DHHC16 constructs for 24h. Proteins were extracted, a total cell extract was analyzed (TCE) and proteins were immunoprecipitated with myc or flag antibodies and subjected to SDS-PAGE, then analyzed by immunoblotting with anti-myc or anti-flag antibodies.

**Figure 2-figure supplement 2.**
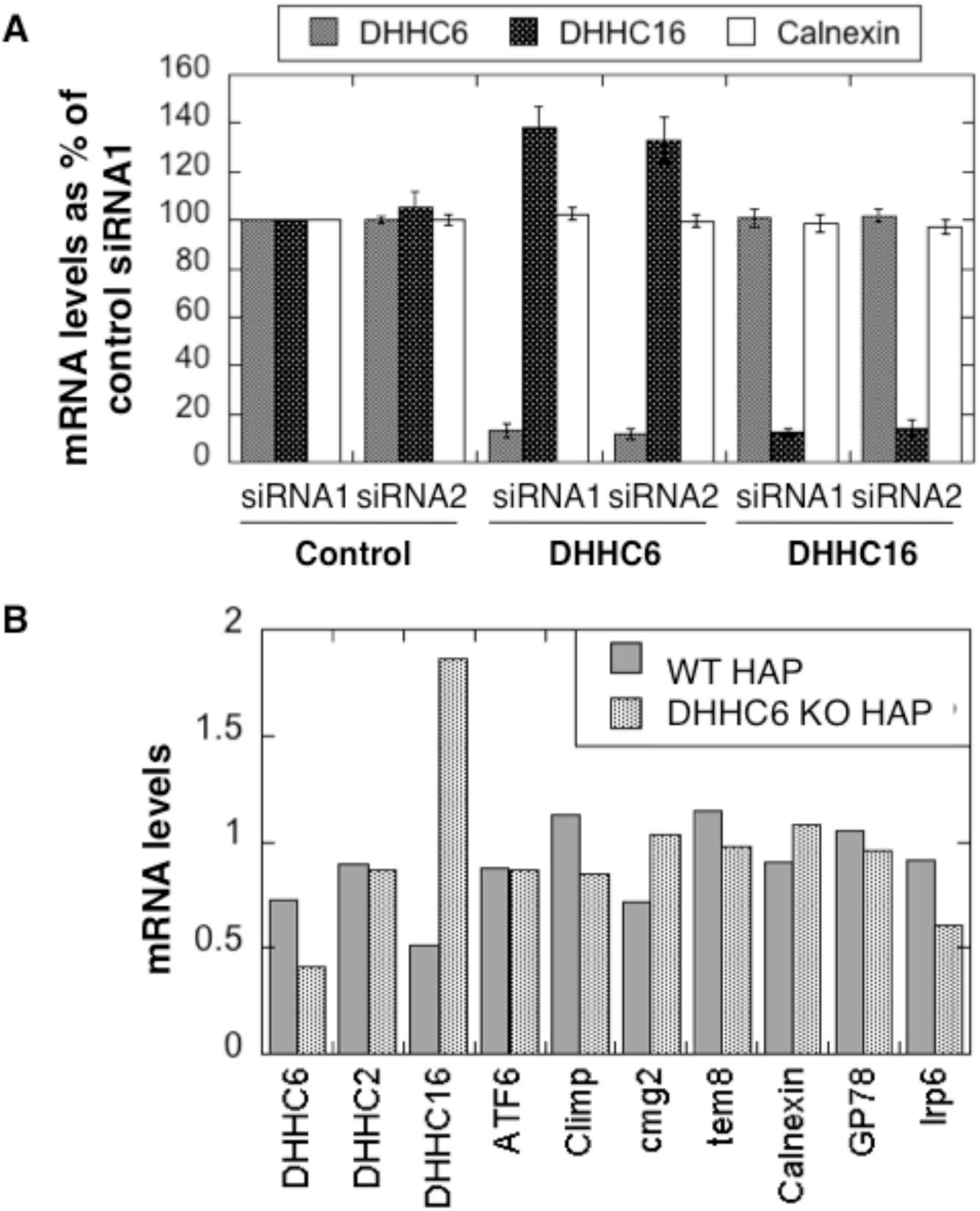
**A.** Silencing of DHHC6 and DHHC16 is efficient. Hela cells were transfected for 72h with either 2 control siRNA or different DHHC siRNA. DHHC6, DHHC16, Calnexin RNA levels were analyzed by quantitative RT-PCR. The histogram shows that silencing was efficient for all DHHC enzymes. N=4. **B.** DHHC16 RNA is increased in DHHC6 KO cells. Annotated RNA levels were analyzed by quantitative RT-PCR in HAP1 cells control or KO for DHHC6.

**Figure 2-figure supplement 3.**
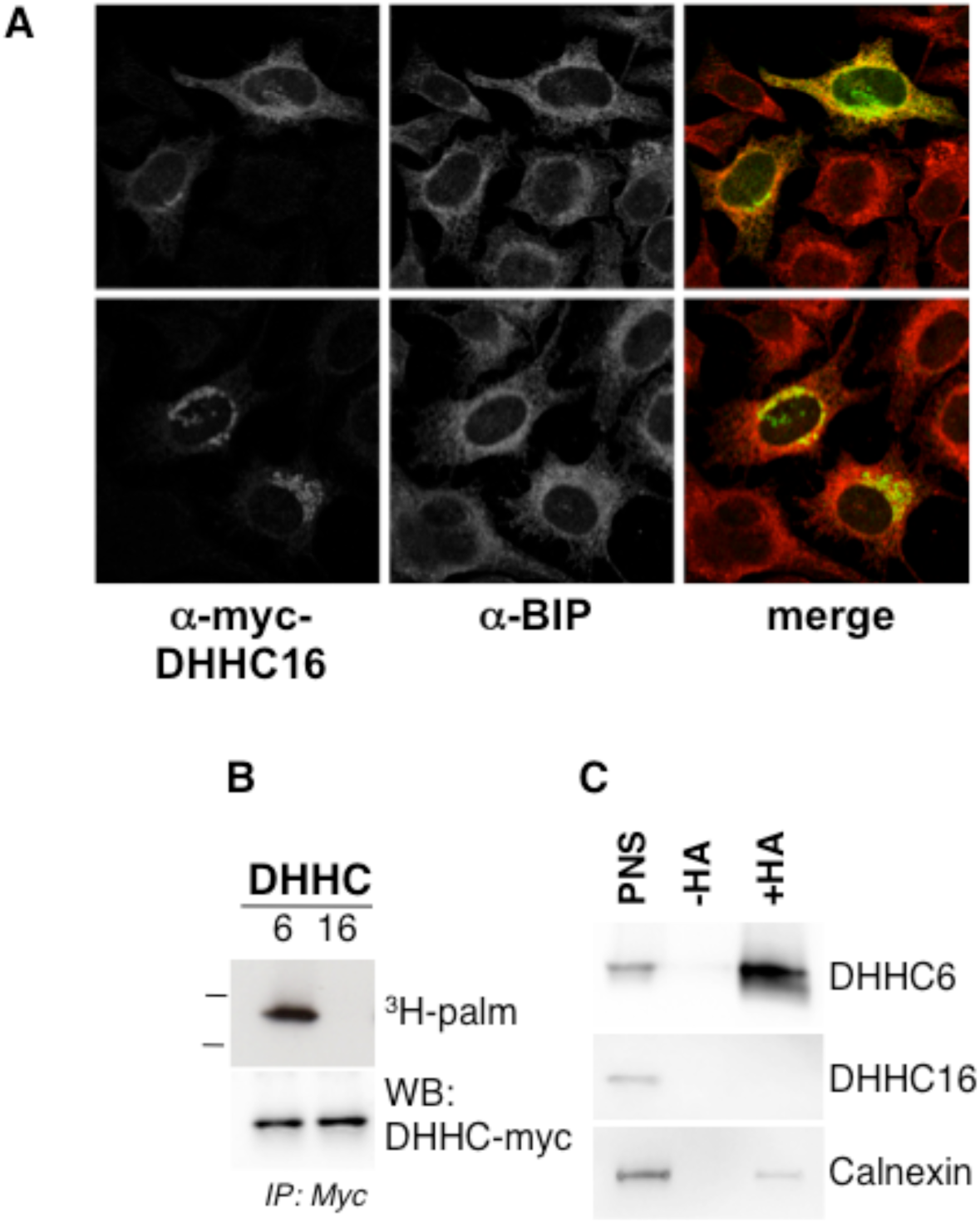
**A.** Immunofluorescence staining of HeLa cells transfected with WT myc-DHHC6 and imaged at high (top lane) or lower expression levels (bottom lane). **B.**DHHC6 is palmitoylated but not DHHC16. Hela cells were transfected with WT myc-tagged DHHC6 or myc-tagged DHHC16 construct for 24h. Cells were then metabolically labeled 2 hours at 37°C with 3H-palmitic acid. Proteins were extracted, immunoprecipitated with myc antibodies and subjected to SDS-PAGE and analyzed by autoradiography, quantified using the Typhoon Imager or by immunoblotting with myc or flag antibodies. **C.** Analysis of protein acylation in Hela cells (Ctrl). Hela were transfected 24h with WT myc-DHHC6 or WT flag-DHHC16 constructs. Cell membranes were recovered by centrifugation and incubated with MMTS and then with hydroxylamine (+HA) or with TRIS (-HA) together with free thiol group binding beads. Eluted fractions were analyzed by immunoblotting with the indicated antibodies. The input fraction (PNS) was loaded as 1/10 this amount.

**Figure 3-figure supplement 1.**
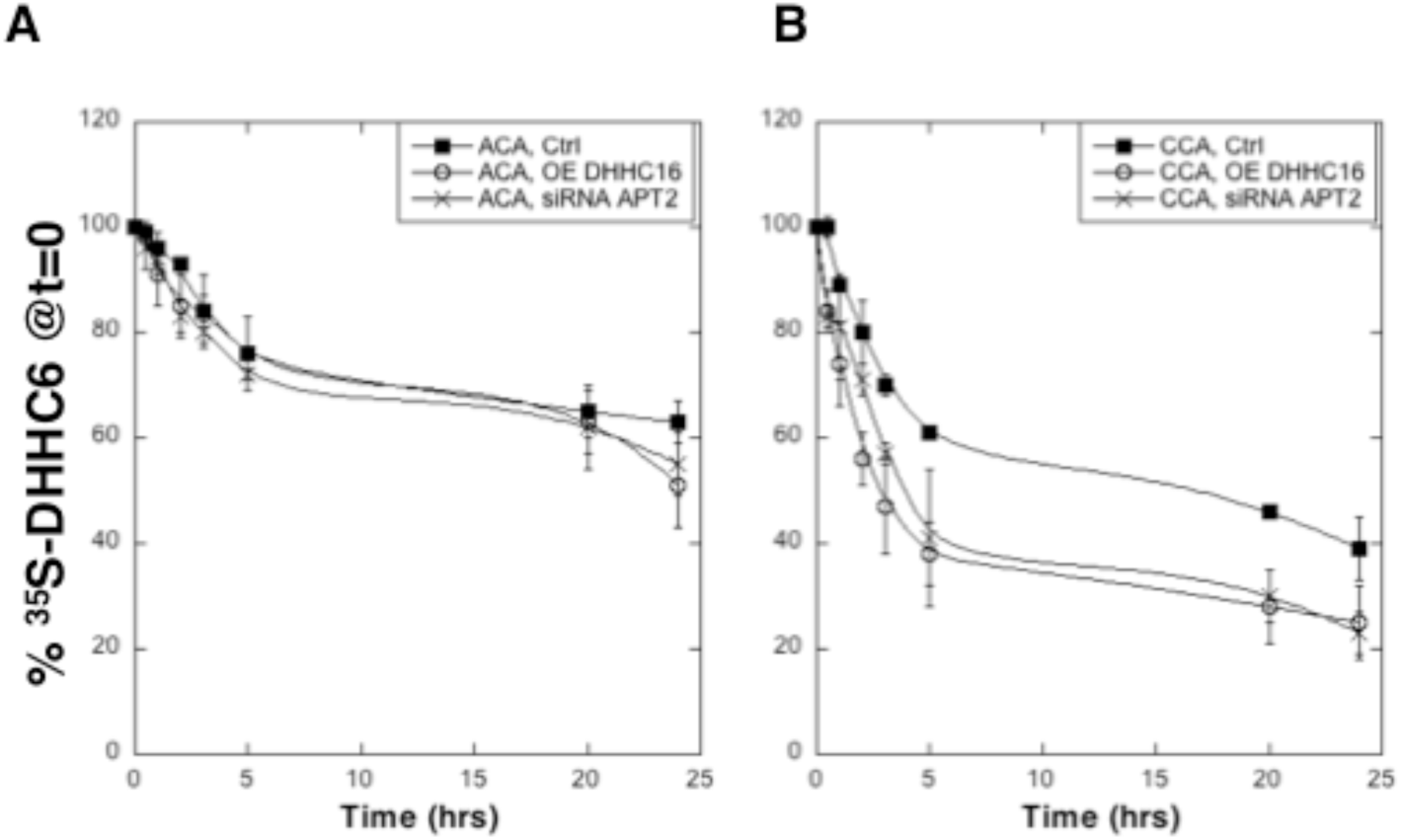
**AB.** Degradation kinetics of DHHC6. Hela cells were transfected with plasmids encoding cysteine mutants myc-DHHC6 constructs with or without WT flag-DHHC16 for 24h after 48h transfection with siRNA APT2 or with control siRNA. Hela cells were incubated 20min pulse with ^35^S-methionin/cysteine at 37°C, washed and further incubated for different times at 37°C in complete medium. DHHC6 were immunoprecipitated and subjected to SDS-PAGE and analyzed by autoradiography, quantified using the Typhoon Imager, and western blotting with anti-myc antibodies. ^35^S-methionin/cysteine incorporation into different DHHC6 constructs were quantified for each times, normalized to protein expression level. The calculated value of ^35^S-methionin/cysteine incorporation into DHHC6 was set to 100% for t=0 after the 20min pulse and all different times of chase with complete medium were expressed relative to this. N=3.

**Figure 5-figure supplement 1.**
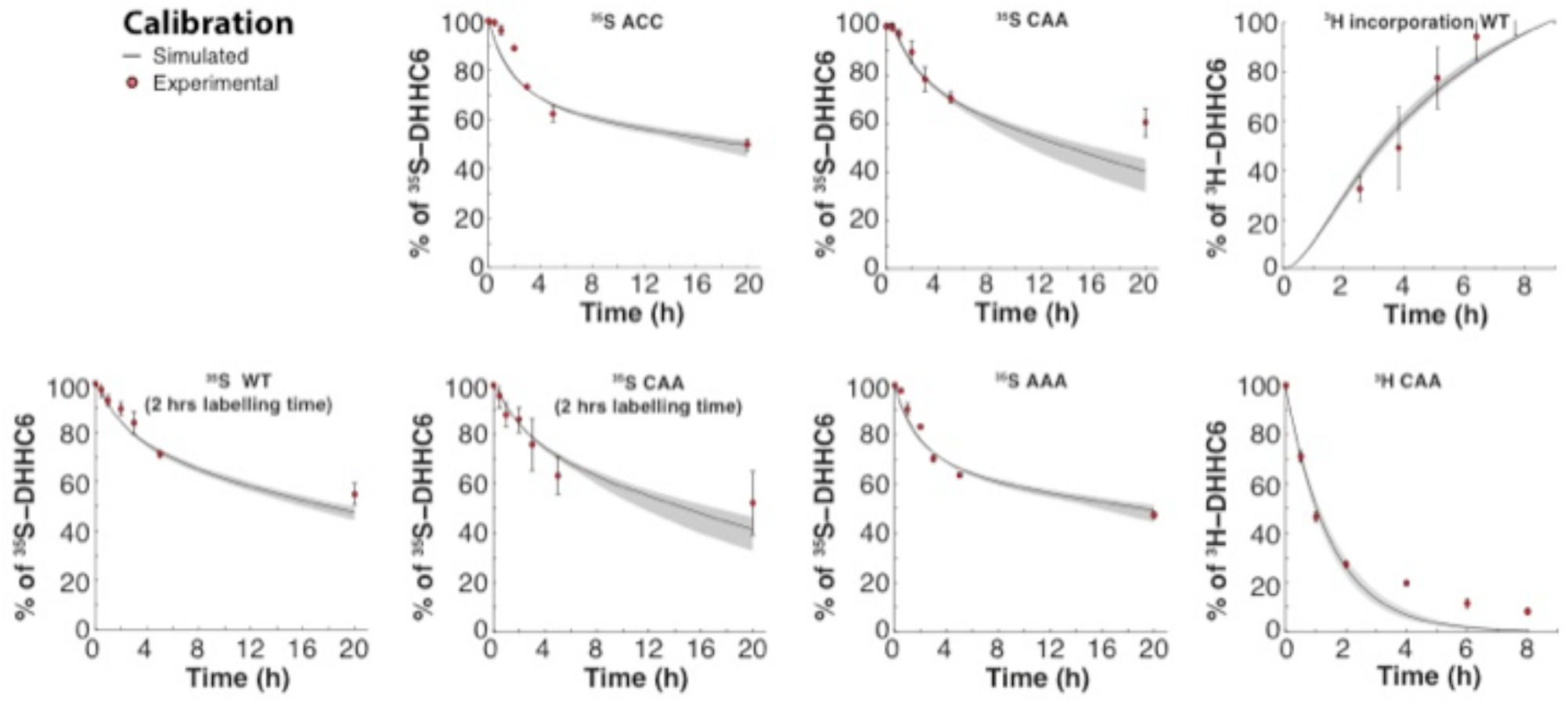
Calibration. Calibration data Results of the GA optimization are plotted on top of the experimental data used as objectives.

**Figure 5-figure supplement 2.**
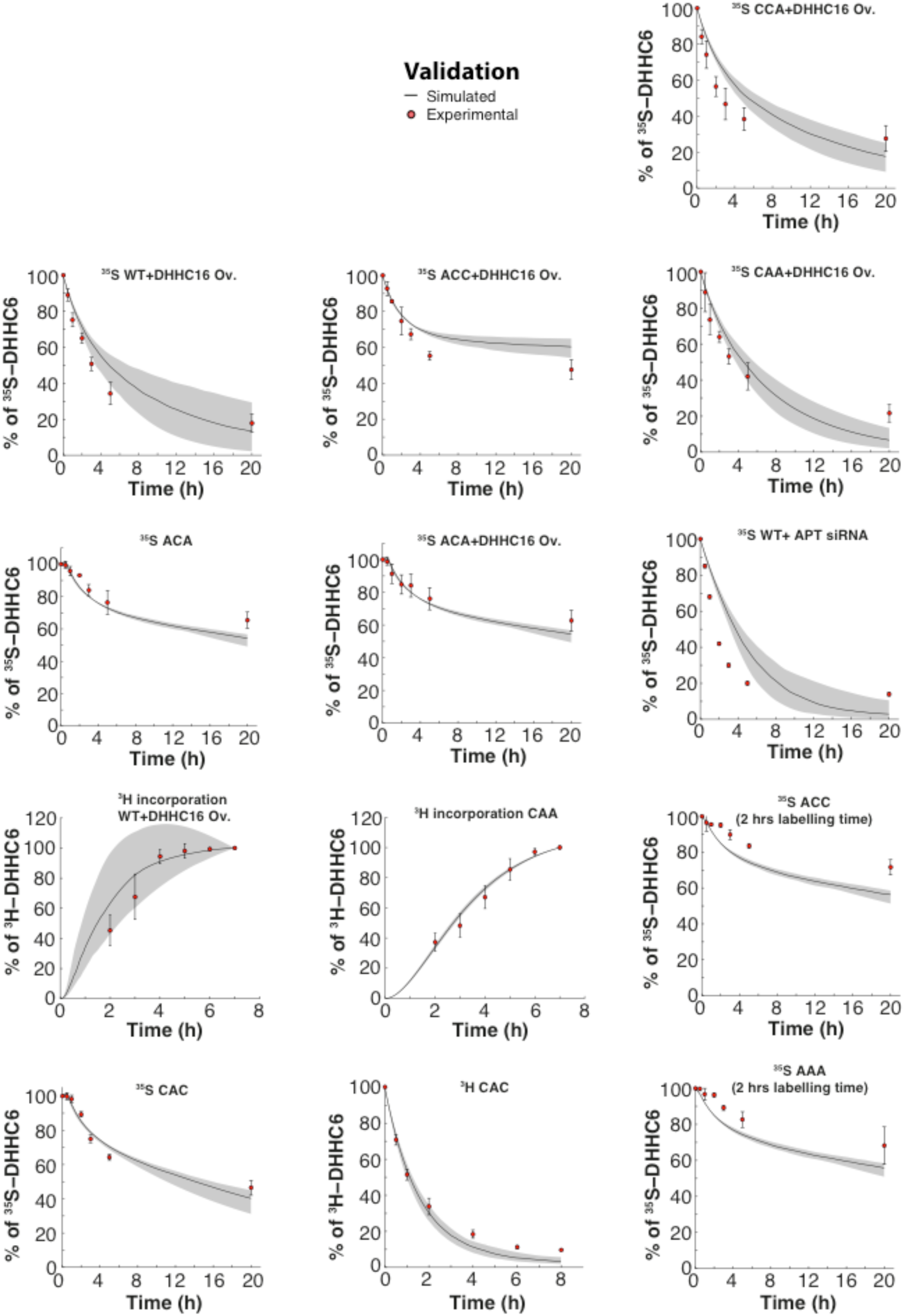
Validation. Results of the GA optimization are plotted on top of the experimental data used as validation.

**Figure 6-figure supplement 1.**
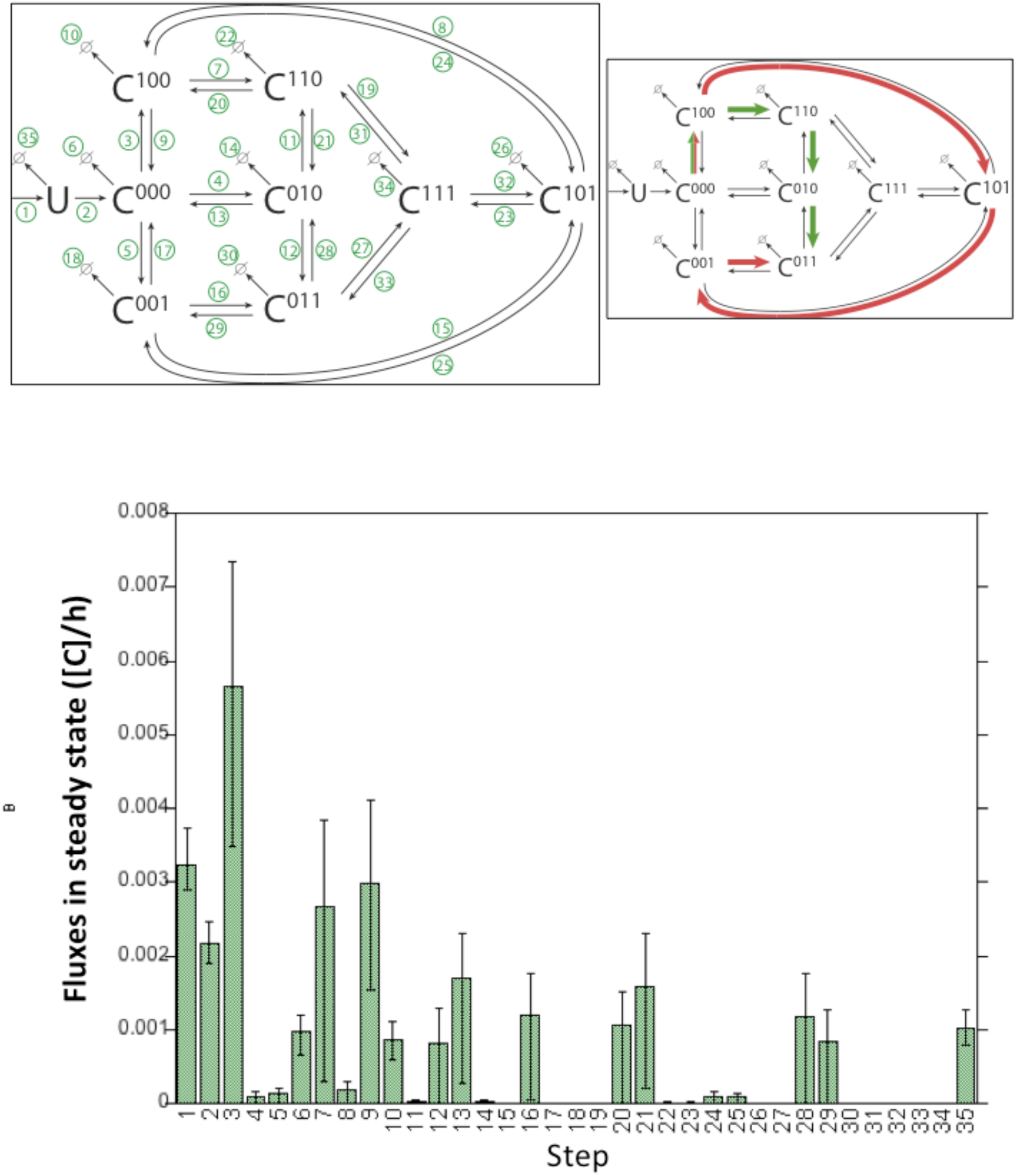
Model fluxes in steady state. The histogram shows the fluxes of the DHHC6 model in steady state under control condition. Lower Panel: Time required by DHHC6 to get palmitoylated for the first time after synthesis in control condition and DHHC6 Overexpression or APT2 silencing conditions. Histograms where obtained by single molecule tracking during stochastic simulations, measuring the time required to get palmitoylated on one of the three available sites.

**Table S1.**
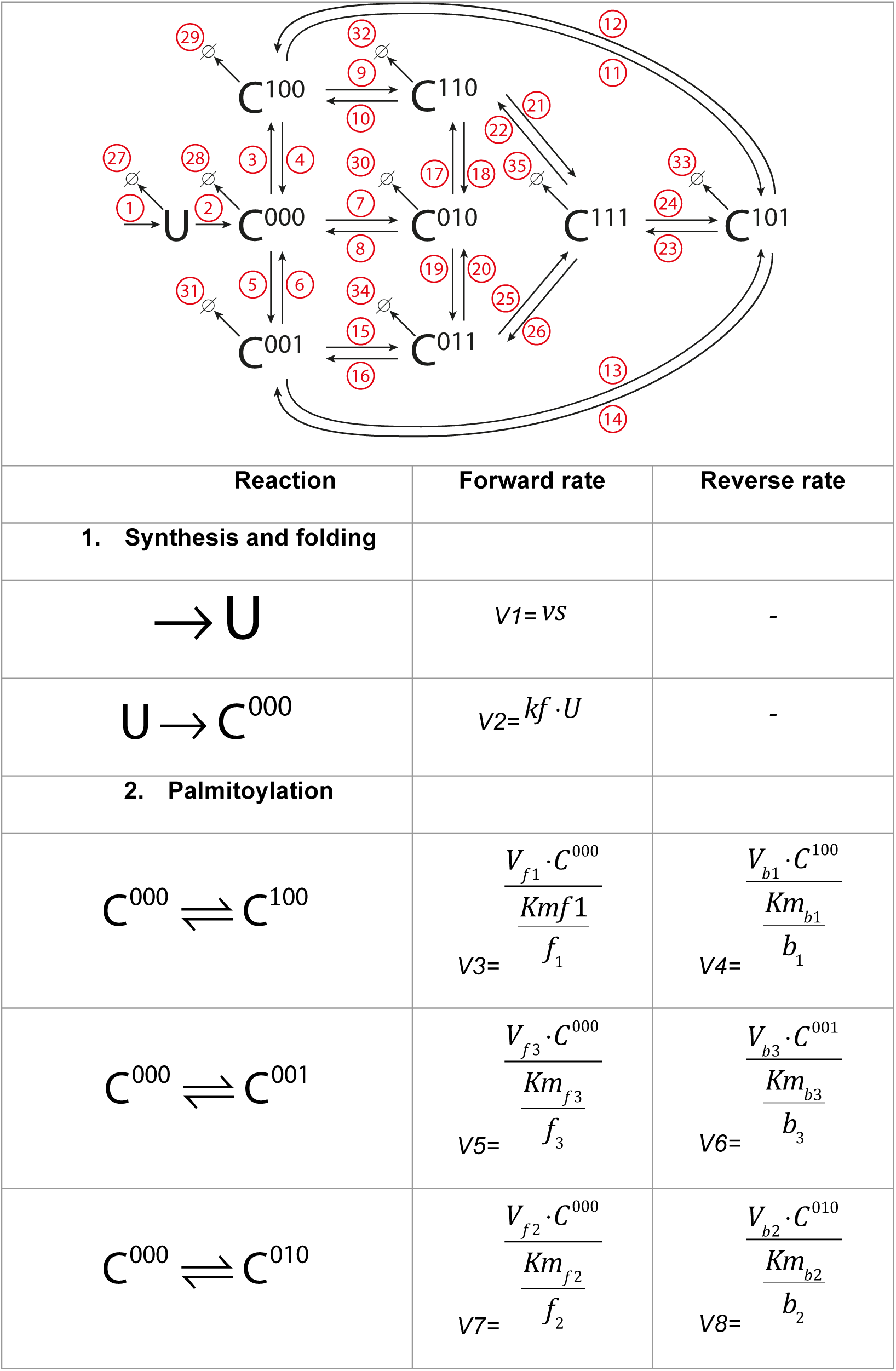
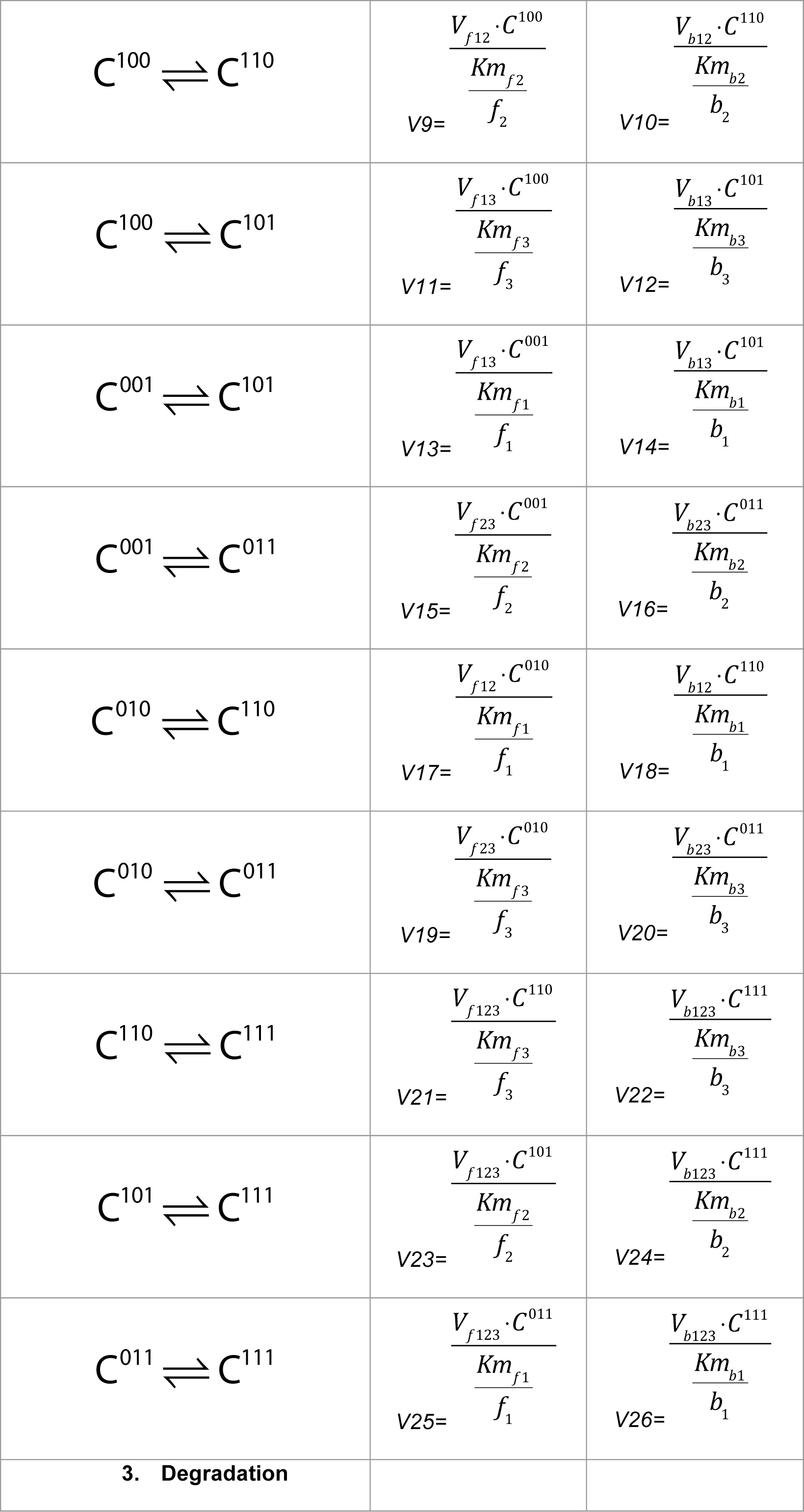
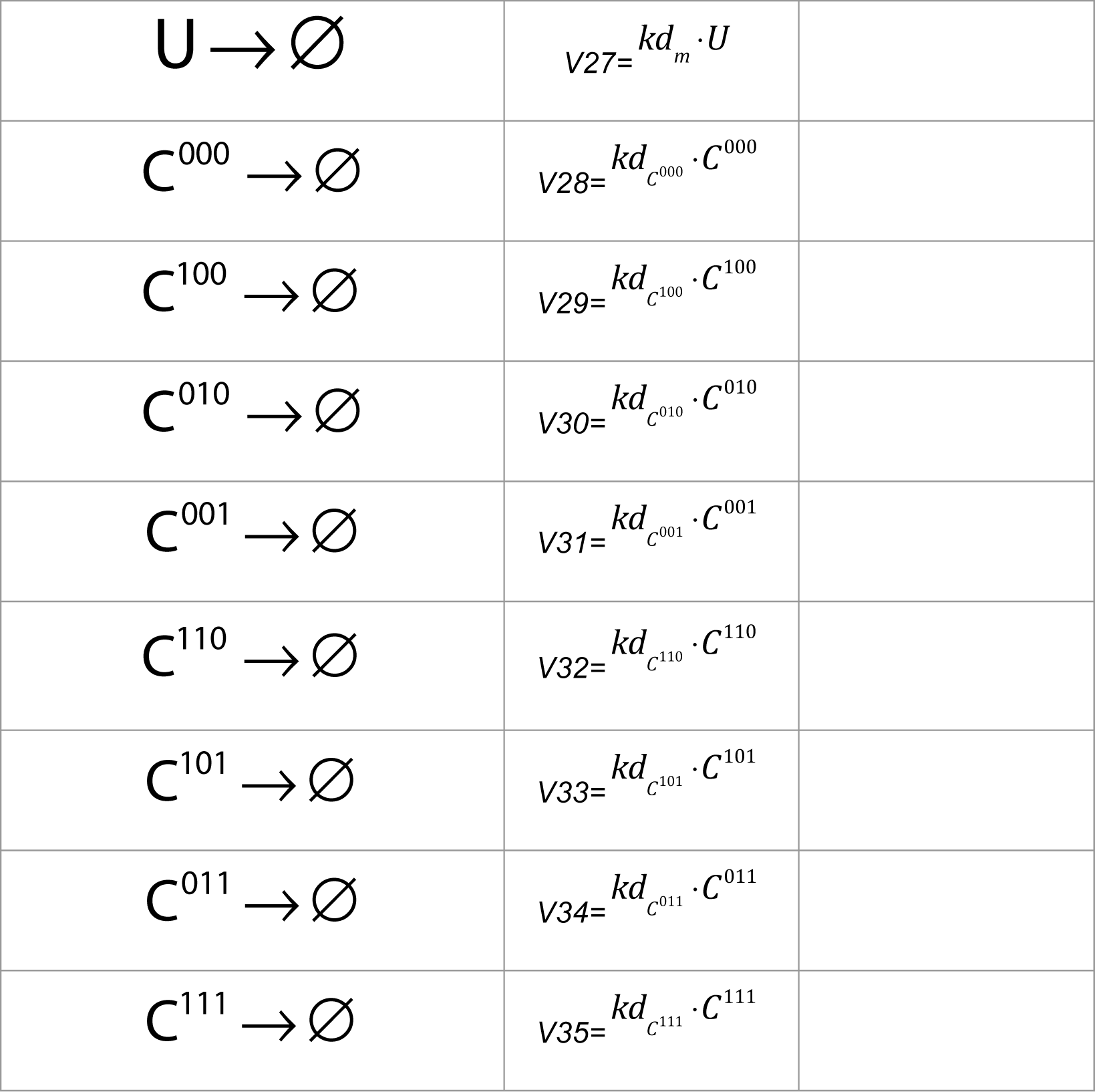
Model reactions. The model of DHHC6 palmitoylation contains 35 different reactions, describing synthesis, folding, degradation, and the enzymatic reactions of DHHC6 palmitoylation/depalmitoylation. In the following table we describe in detail how the rates for those reactions are calculated.

Where the competition terms *f* and *b* in S1 Table are defined as:

|  |  |
| --- | --- |
| $f_1$ | $1 + \frac{C^{000}}{Km_{f1}} + \frac{C^{000}}{Km_{f2}} + \frac{C^{000}}{Km_{f3}} + \frac{C^{100}}{Km_{f2}} + \frac{C^{100}}{Km_{f3}} + \frac{C^{010}}{Km_{f1}} + \frac{C^{010}}{Km_{f3}} + \frac{C^{001}}{Km_{f1}} + \frac{C^{001}}{Km_{f2}} + \frac{C^{110}}{Km_{f3}} + \frac{C^{101}}{Km_{f2}} + \frac{C^{011}}{Km_{f1}} + \frac{DHHC_{16t}}{Km_{f1}}$ |
| $f_2$ | $1 + \frac{C^{000}}{Km_{f1}} + \frac{C^{000}}{Km_{f2}} + \frac{C^{000}}{Km_{f3}} + \frac{C^{100}}{Km_{f2}} + \frac{C^{100}}{Km_{f3}} + \frac{C^{010}}{Km_{f1}} + \frac{C^{010}}{Km_{f3}} + \frac{C^{001}}{Km_{f1}} + \frac{C^{001}}{Km_{f2}} + \frac{C^{110}}{Km_{f3}} + \frac{C^{101}}{Km_{f2}} + \frac{C^{011}}{Km_{f1}} + \frac{DHHC_{16t}}{Km_{f2}}$ |
| $f_3$ | $1 + \frac{C^{000}}{Km_{f1}} + \frac{C^{000}}{Km_{f2}} + \frac{C^{000}}{Km_{f3}} + \frac{C^{100}}{Km_{f2}} + \frac{C^{100}}{Km_{f3}} + \frac{C^{010}}{Km_{f1}} + \frac{C^{010}}{Km_{f3}} + \frac{C^{001}}{Km_{f1}} + \frac{C^{001}}{Km_{f2}} + \frac{C^{110}}{Km_{f3}} + \frac{C^{101}}{Km_{f2}} + \frac{C^{011}}{Km_{f1}} + \frac{DHHC_{16t}}{Km_{f3}}$ |
| $b_1$ | $1 + \frac{C^{111}}{Km_{b3}} + \frac{C^{111}}{Km_{b2}} + \frac{C^{111}}{Km_{b1}} + \frac{C^{100}}{Km_{b1}} + \frac{C^{010}}{Km_{b2}} + \frac{C^{001}}{Km_{b3}} + \frac{C^{110}}{Km_{b1}} + \frac{C^{110}}{Km_{b2}} + \frac{C^{101}}{Km_{b1}} + \frac{C^{101}}{Km_{b3}} + \frac{C^{011}}{Km_{b3}} + \frac{C^{011}}{Km_{b2}} + \frac{APT_{2t}}{Km_{b1}}$ |
| $b_2$ | $1 + \frac{C^{111}}{Km_{b3}} + \frac{C^{111}}{Km_{b2}} + \frac{C^{111}}{Km_{b1}} + \frac{C^{100}}{Km_{b1}} + \frac{C^{010}}{Km_{b2}} + \frac{C^{001}}{Km_{b3}} + \frac{C^{110}}{Km_{b1}} + \frac{C^{110}}{Km_{b2}} + \frac{C^{101}}{Km_{b1}} + \frac{C^{101}}{Km_{b3}} + \frac{C^{011}}{Km_{b3}} + \frac{C^{011}}{Km_{b2}} + \frac{APT_{2t}}{Km_{b2}}$ |
| $b_3$ | $1 + \frac{C^{111}}{Km_{b3}} + \frac{C^{111}}{Km_{b2}} + \frac{C^{111}}{Km_{b1}} + \frac{C^{100}}{Km_{b1}} + \frac{C^{010}}{Km_{b2}} + \frac{C^{001}}{Km_{b3}} + \frac{C^{110}}{Km_{b1}} + \frac{C^{110}}{Km_{b2}} + \frac{C^{101}}{Km_{b1}} + \frac{C^{101}}{Km_{b3}} + \frac{C^{011}}{Km_{b3}} + \frac{C^{011}}{Km_{b2}} + \frac{APT_{2t}}{Km_{b3}}$ |

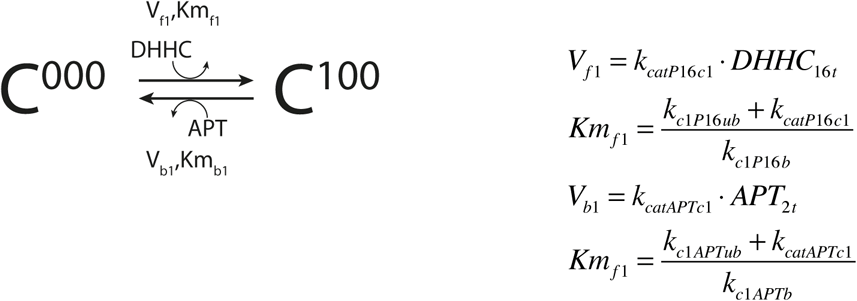

where *k*_*catP16c1*_ is the catalytic constant for DHHC16 when catalyzing palmitoylation of C^100^ on DHHC6, while *k*_*catAPTc1*_ is the catalytic constant for APT2 for the same site. *k*_*c*1*P*16 *b*_ and *k*_*c*1*P*16*ub*_ are the binding and unbinding rate constants of DHHC6 to DHHC16 on site C^100^ respectively. *K*_*c1APTb*_ and *k*_*c1APTub*_ *k*_c1APTub_ are the binding and unbinding rate constants of DHHC6 to APT1 on site C^100^ respectively. In our model we assume that the binding and unbinding rate to DHHC6 and APT can be different for each state of the model.

**Table S2.**
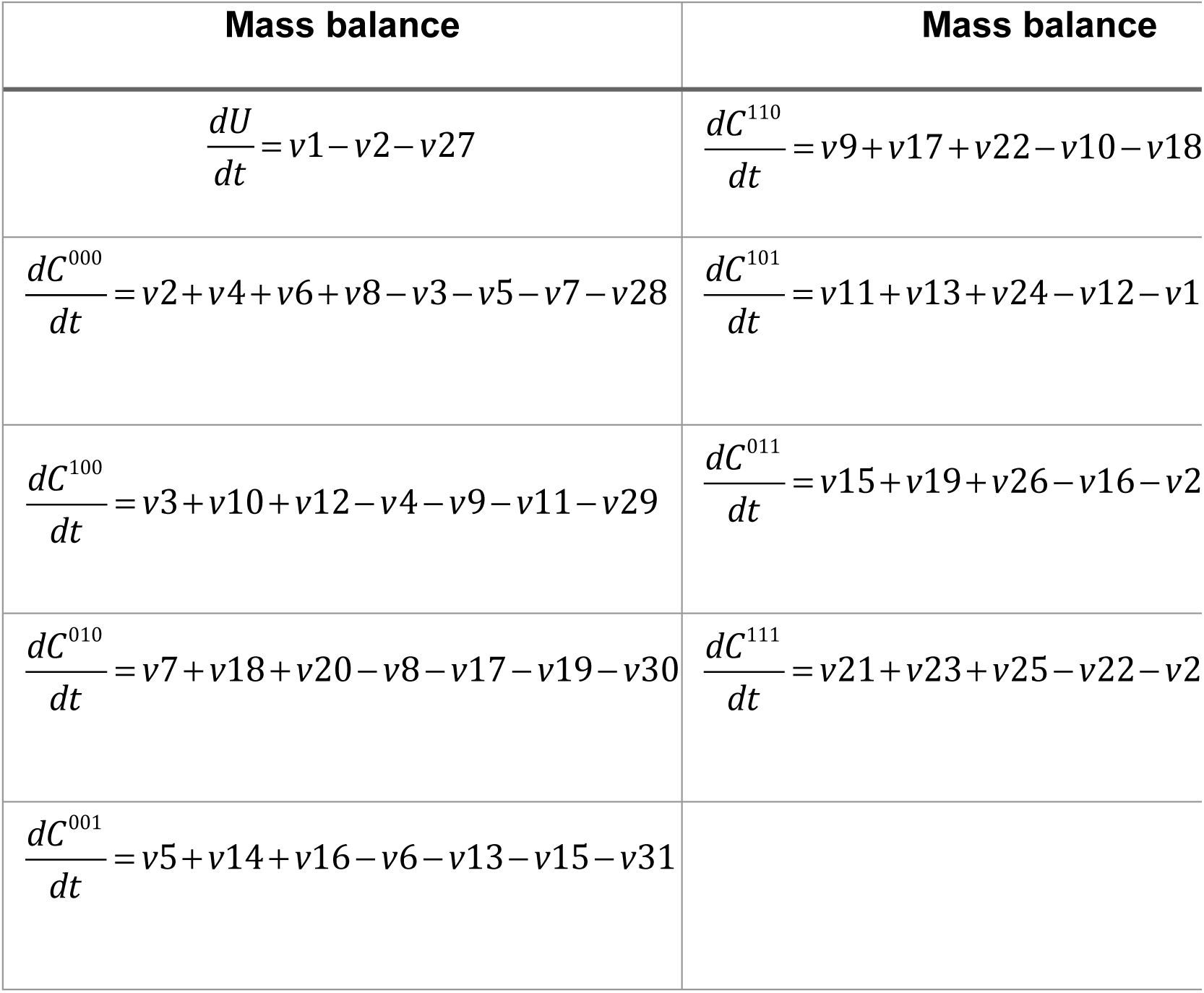
Mass balance equations. The following table describes the mass balance for each of the species of DHHC6 model. The rates of the mass balance each state are described in detail in S1 Table.

**Table S3.** Model parameters. The output of GA is a set of optimal solutions, where a solution is a complete set of parameter needed to perform model simulations. From this set we extracted a sub-set of 152 solutions that obtained a GA score better than a set threshold for each objective. During the analysis the model was simulated for each set of parameters of the sub-set. We then reported in this paper the mean of the outputs along with the 1^st^ and 3^rd^ quartile of their distribution.

| Parameter | Median Value | Range |
| --- | --- | --- |
| $v_s$ [C]<sup>a</sup>/[T]<sup>b</sup> | $3.00 \cdot 10^{-3}$ | $[2.59 \cdot 10^{-3} - 3.31 \cdot 10^{-3}]$ |
| $k_f$ 1/[T] | $4.20 \cdot 10^{-1}$ | $[2.98 \cdot 10^{-1} - 6.69 \cdot 10^{-1}]$ |
| $V_{f1}$ [C]/[T] | 6.03 | [4.54-6.92] |
| $Km_{f1}$ [C] | $2.02 \cdot 10^{-2}$ | $[1.62 \cdot 10^{-2} - 2.48 \cdot 10^{-2}]$ |
| $V_{f2}$ [C]/[T] | $4.21 \cdot 10^{-4}$ | $[1.18 \cdot 10^{-4} - 9.89 \cdot 10^{-4}]$ |
| $Km_{f2}$ [C] | $1.33 \cdot 10^{-5}$ | $[9.40 \cdot 10^{-6} - 2.16 \cdot 10^{-5}]$ |
| $V_{f3}$ [C]/[T] | $8.07 \cdot 10^{-2}$ | $[4.30 \cdot 10^{-2} - 1.49 \cdot 10^{-1}]$ |
| $Km_{f3}$ [C] | $1.06 \cdot 10^{-2}$ | $[8.21 \cdot 10^{-3} - 1.22 \cdot 10^{-2}]$ |
| $V_{b3}$ [C]/[T] | $8.12 \cdot 10^{-3}$ | $[5.07 \cdot 10^{-3} - 1.13 \cdot 10^{-2}]$ |
| $Km_{b3}$ [C] | $1.63 \cdot 10^{-3}$ | $[7.99 \cdot 10^{-4} - 3.22 \cdot 10^{-3}]$ |
| $V_{b2}$ [C]/[T] | $5.86 \cdot 10^{-2}$ | $[4.52 \cdot 10^{-2} - 7.44 \cdot 10^{-2}]$ |
| $Km_{b2}$ [C] | $1.74 \cdot 10^{-2}$ | $[1.18 \cdot 10^{-2} - 2.29 \cdot 10^{-2}]$ |
| $V_{b1}$ [C]/[T] | $1.18 \cdot 10^{-1}$ | $[9.04 \cdot 10^{-2} - 1.42 \cdot 10^{-1}]$ |
| $Km_{b1}$ [C] | $3.71 \cdot 10^{-3}$ | $[2.27 \cdot 10^{-3} - 6.13 \cdot 10^{-3}]$ |
| $kd_m$ 1/[T] | $2.56 \cdot 10^{-1}$ | $[2.16 \cdot 10^{-1} - 3.50 \cdot 10^{-1}]$ |
| $kd_{c^{000}}$ 1/[T] | $1.80 \cdot 10^{-2}$ | $[1.47 \cdot 10^{-2} - 2.72 \cdot 10^{-2}]$ |
| $kd_{c^{100}}$ 1/[T] | $1.33 \cdot 10^{-1}$ | $[8.81 \cdot 10^{-2} - 1.97 \cdot 10^{-1}]$ |
| $kd_{c^{010}}$ 1/[T] | $4.82 \cdot 10^{-3}$ | $[1.58 \cdot 10^{-3} - 1.32 \cdot 10^{-2}]$ |
| $kd_{c^{001}}$ 1/[T] | $3.89 \cdot 10^{-2}$ | $[2.07 \cdot 10^{-2} - 6.04 \cdot 10^{-2}]$ |
| $kd_{c^{110}}$ 1/[T] | $1.28 \cdot 10^{-1}$ | $[3.66 \cdot 10^{-2} - 2.50 \cdot 10^{-1}]$ |
| $kd_{c^{101}}$ 1/[T] | $5.59 \cdot 10^{-2}$ | $[2.74 \cdot 10^{-2} - 2.41 \cdot 10^{-1}]$ |
| $kd_{c^{011}}$ 1/[T] | $3.42 \cdot 10^{-4}$ | $[6.46 \cdot 10^{-5} - 1.36 \cdot 10^{-3}]$ |
| $kd_{c^{111}}$ 1/[T] | 2.32 | $[7.14 \cdot 10^{-1} - 7.31]$ |
| $V_{f12}$ [C]/[T] | $6.43 \cdot 10^{-2}$ | $[3.36 \cdot 10^{-2} - 1.07 \cdot 10^{-1}]$ |
| $V_{f13}$ [C]/[T] | $6.04 \cdot 10^{-1}$ | $[1.54 \cdot 10^{-1} - 1.36]$ |
| $V_{f23}$ [C]/[T] | 2.89 | [1.58-4.74] |
| $V_{f123}$ [C]/[T] | $5.61 \cdot 10^{-1}$ | $[1.37 \cdot 10^{-1} - 2.43]$ |
| $V_{b12}$ [C]/[T] | 1.83 | $[7.00 \cdot 10^{-1} - 5.64]$ |
| $V_{b13}$ [C]/[T] | 1.40 | $[5.01 \cdot 10^{-1} - 5.28]$ |
| $V_{b23}$ [C]/[T] | $9.49 \cdot 10^{-2}$ | $[5.60 \cdot 10^{-2} - 1.54 \cdot 10^{-1}]$ |
| $V_{b123}$ [C]/[T] | $9.35 \cdot 10^{-1}$ | $[2.09 \cdot 10^{-1} - 4.19]$ |
| $DHHC_{16t}$ [C] | $1.73 \cdot 10^{-1}$ | $[1.05 \cdot 10^{-1} - 3.10 \cdot 10^{-1}]$ |
| $APT_{2t}$ [C] | $1.85 \cdot 10^{-1}$ | $[1.42 \cdot 10^{-1} - 2.20 \cdot 10^{-1}]$ |

**Table S4.**
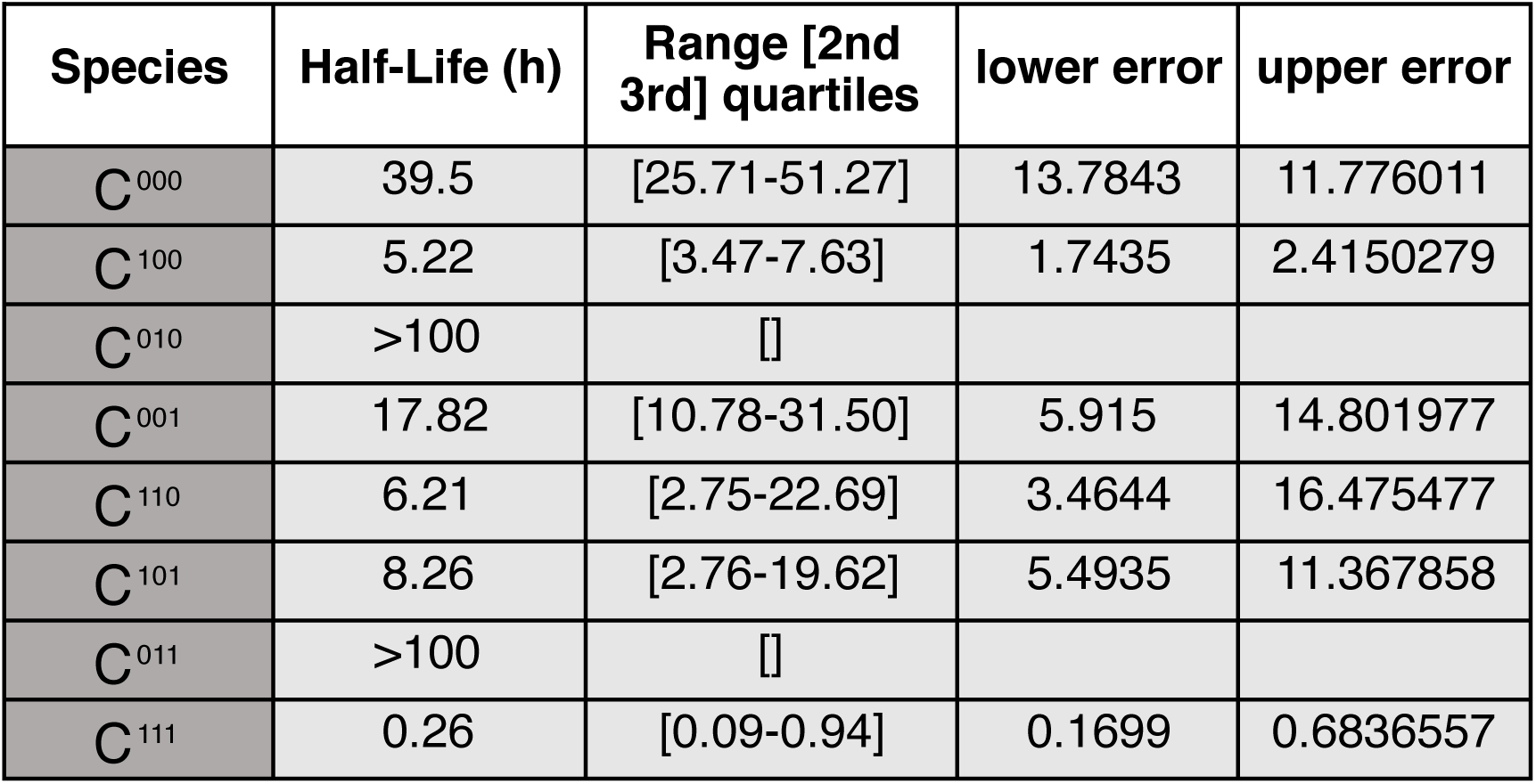
Half-life of DHHC6 in different palmitoylation states. The half-life was estimated from the decay rate constant obtained through parameter estimation. The half-life is calculate as: ln(2)/kd.

**Table S5.** Total amount of protein in steady state relative to WT for WT and CAA mutant in different conditions. The table shows the total protein in steady state in the model relative to the abundance of DHHC6 observed in steady state in WT conditions. Simulations are performed for WT and CAA mutant under control conditions, after overexpression of DHHC16 and after silencing of APT2.

| Protein | Condition | Amount (relative to WT control) | Range [2nd 3rd] quartiles | lower error | upper error |
| --- | --- | --- | --- | --- | --- |
| WT | Control | 100% | [70-112] | 30 | 12 |
| WT | OE DHHC16 | 84% | [27-104] | 47 | 20 |
| WT | siRNA APT2 | 28% | [14-34.54] | 14 | 6.52 |
| CAA | Control | 73% | [43-91] | 30 | 18 |
| CAA | OE DHHC16 | 31% | [19.60-41.50] | 12 | 10.5 |
| CAA | siRNA APT2 | 31% | [20-38] | 11 | 7 |

**Table S6.**
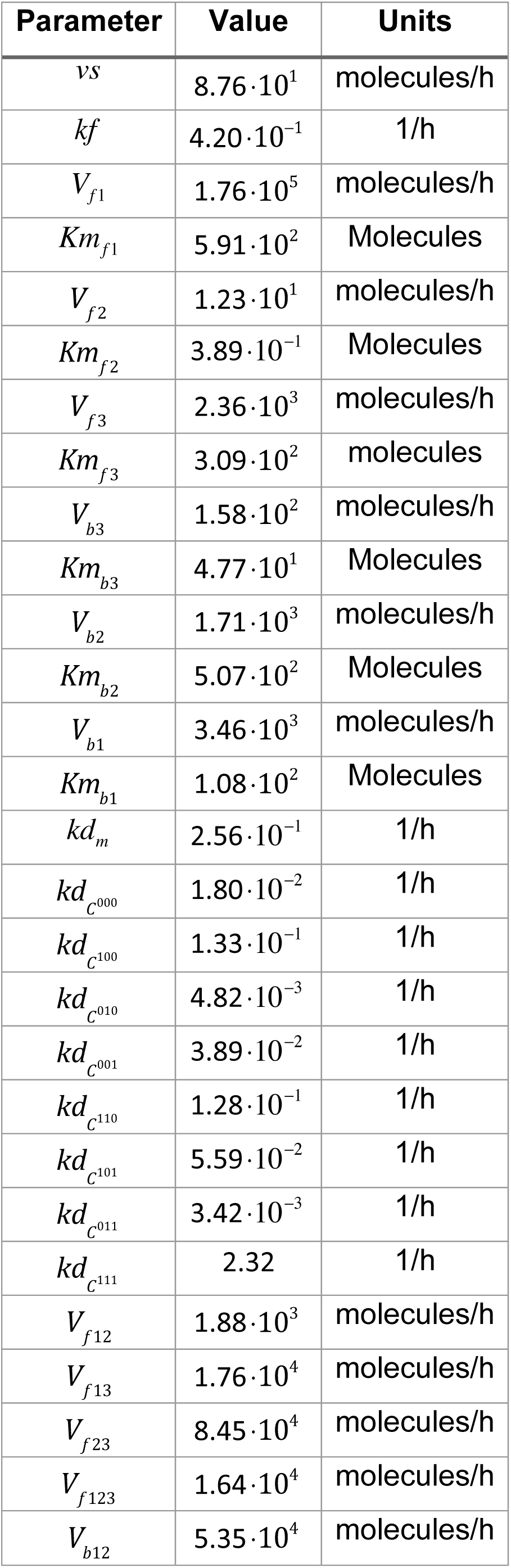
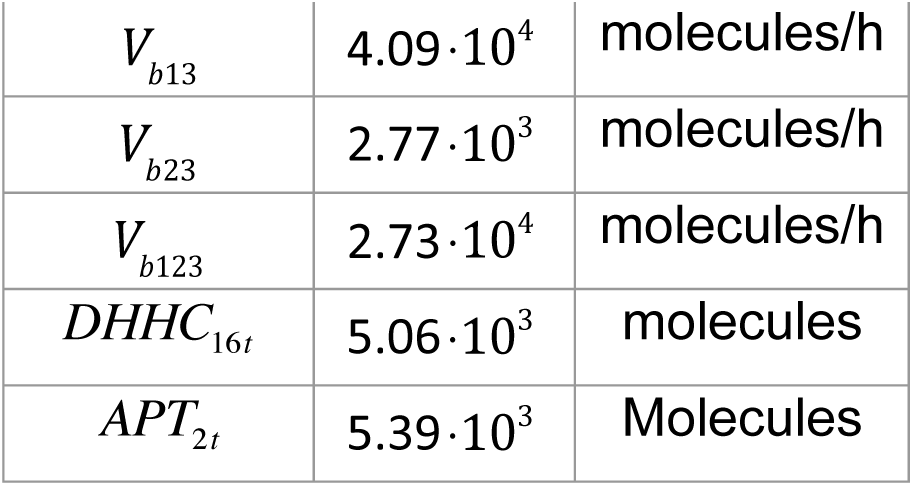
Parameters used for stochastic simulations. The following parameters were obtained through the conversion of the deterministic parameters estimated by the GA (see {Dallavilla, 2016 #211}).

**Table S7.**
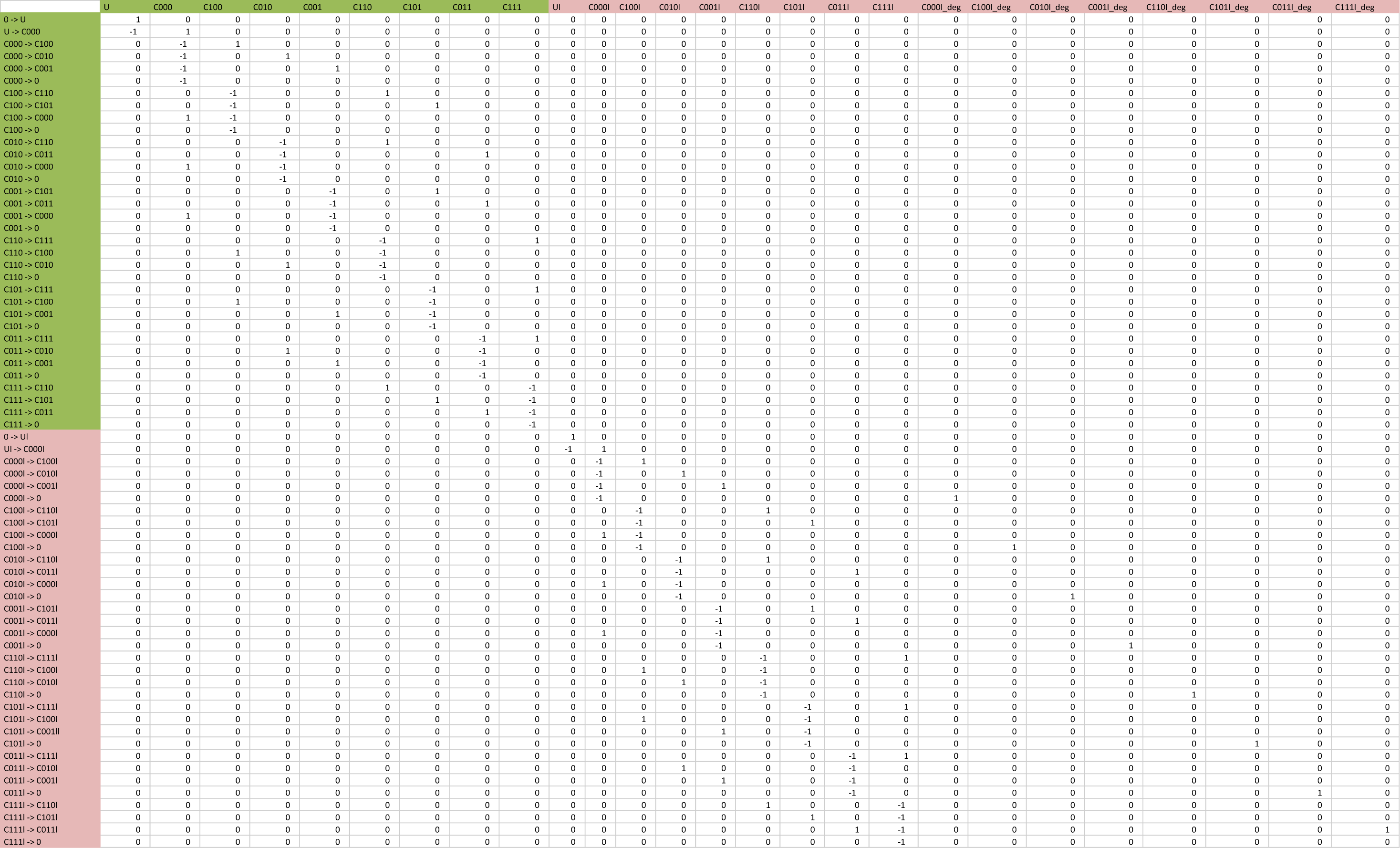
Stoichiometric matrix used for stochastic simulations. In the following table we define the stoichiometry and the directionality of the reactions of the model. Each reaction has a directionality that define which are the reagents and which are the products. In this table each line represents a model reaction, while in the columns we find all the states of the model. For each reaction the states of the model that take part to the reaction as reagents are marked with -1, while the states that participate as products are marked with 1. The matrix that is formed in this way allow to attribute the correct directionality to model reactions during the calculation of the mass balance for each state of the model.

**Table S8.**
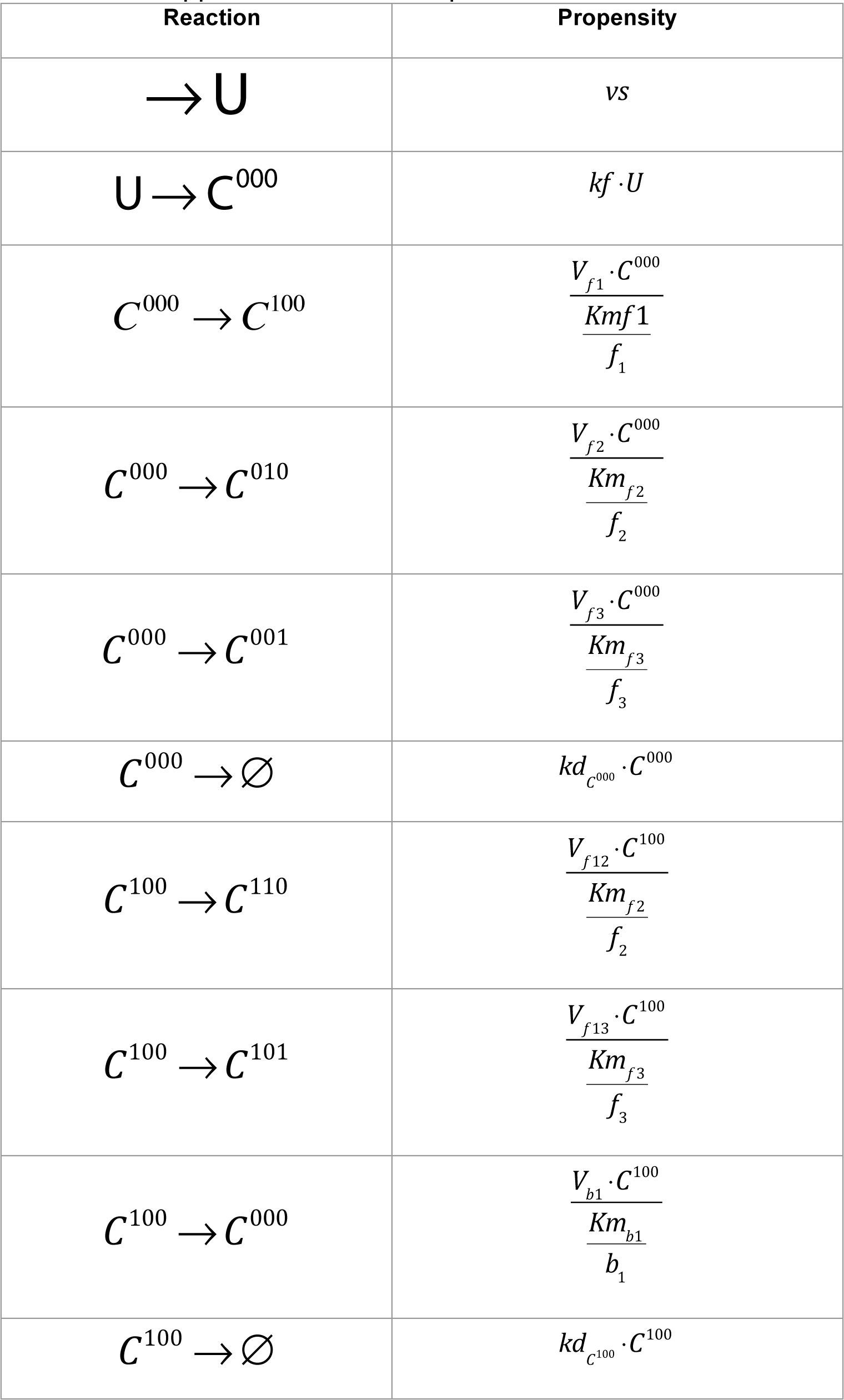
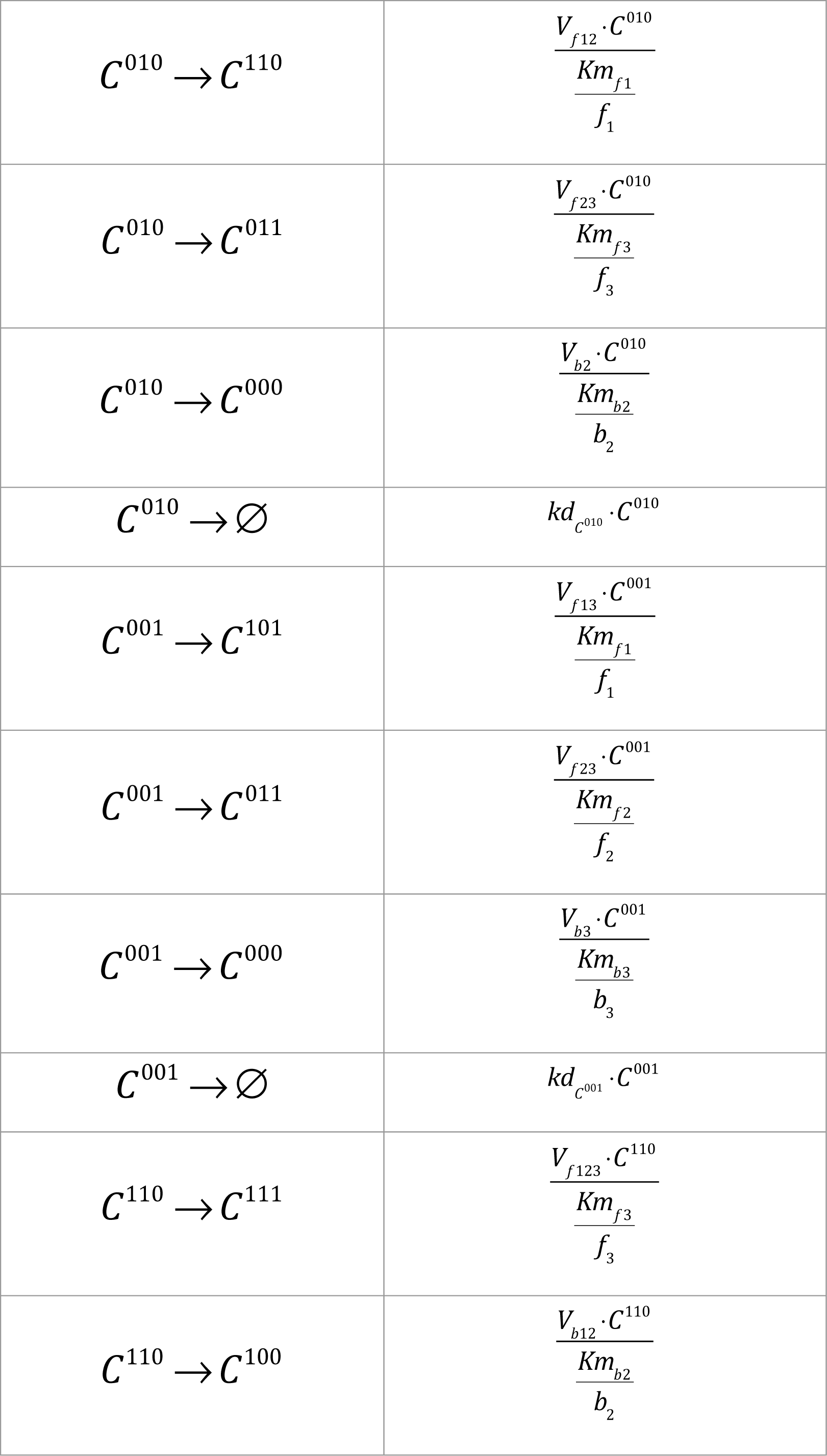
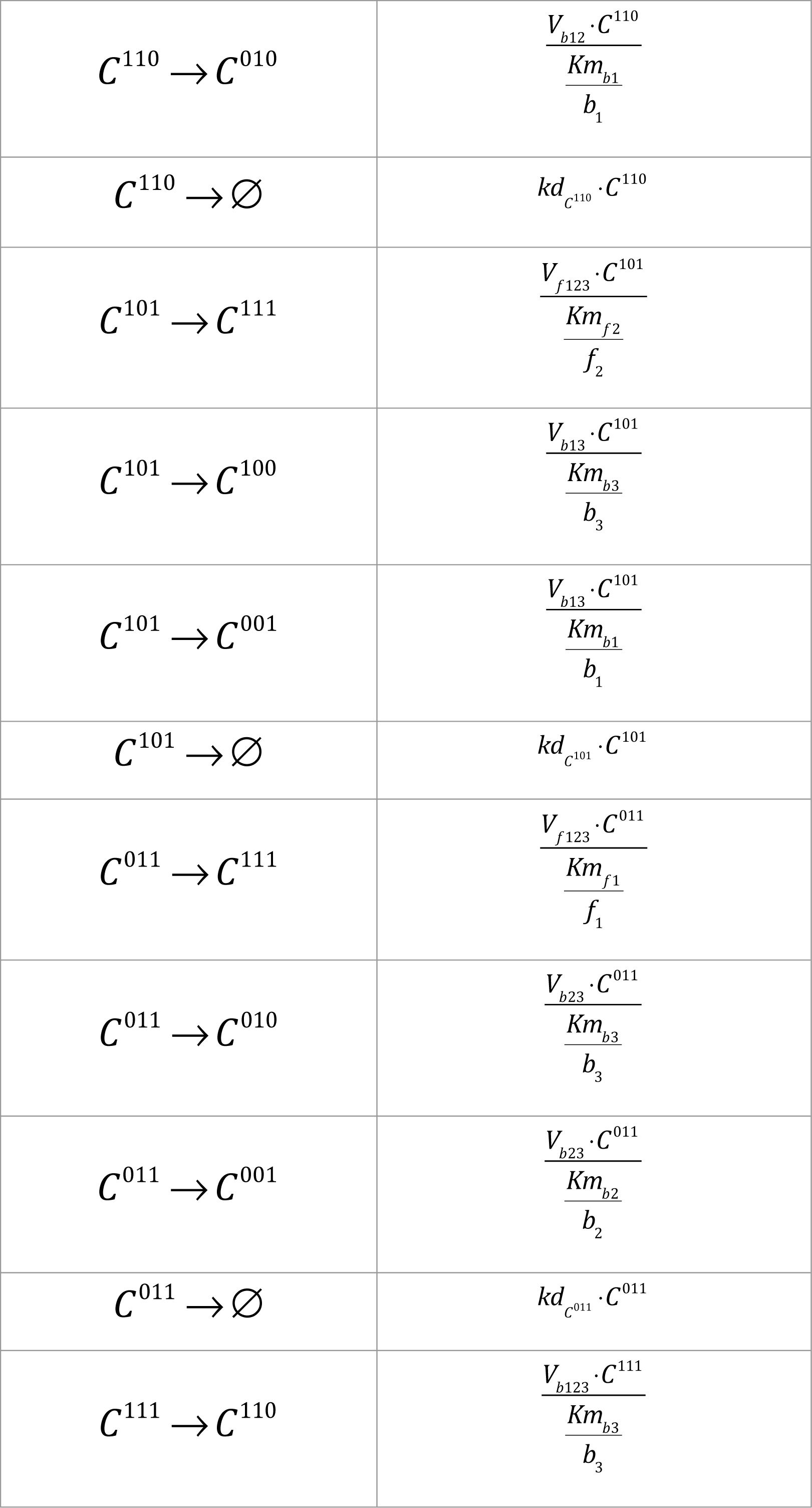
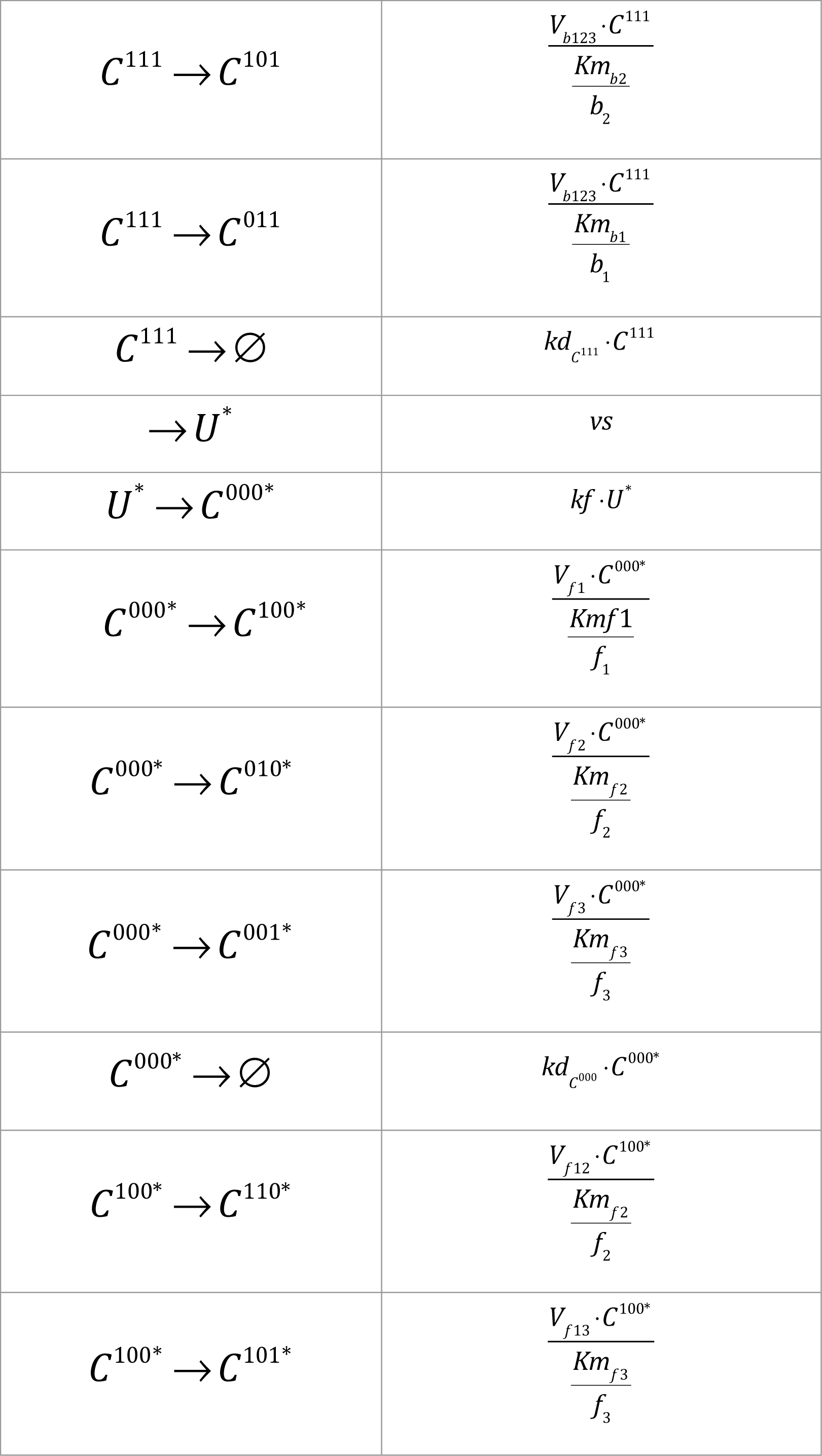
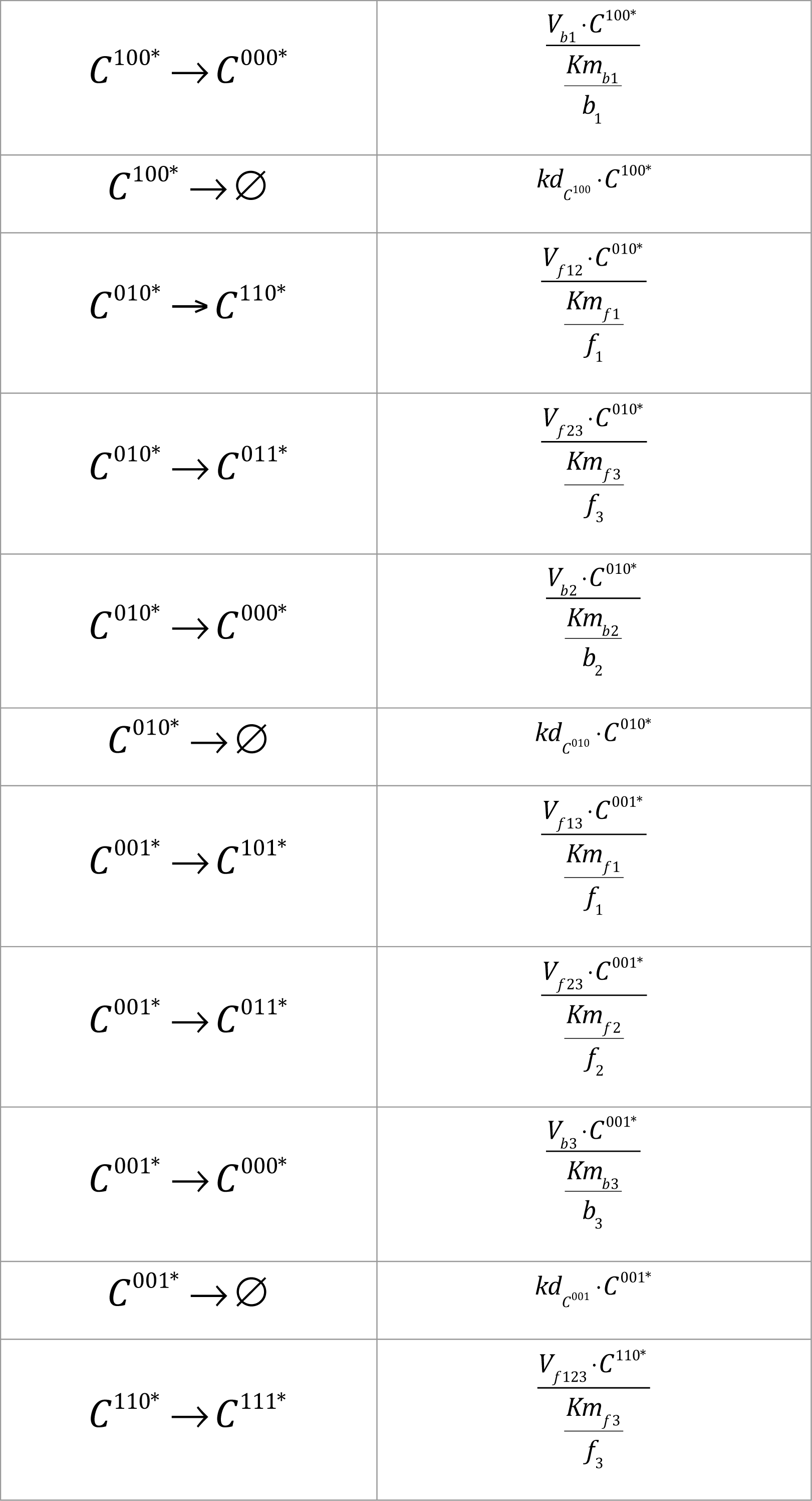
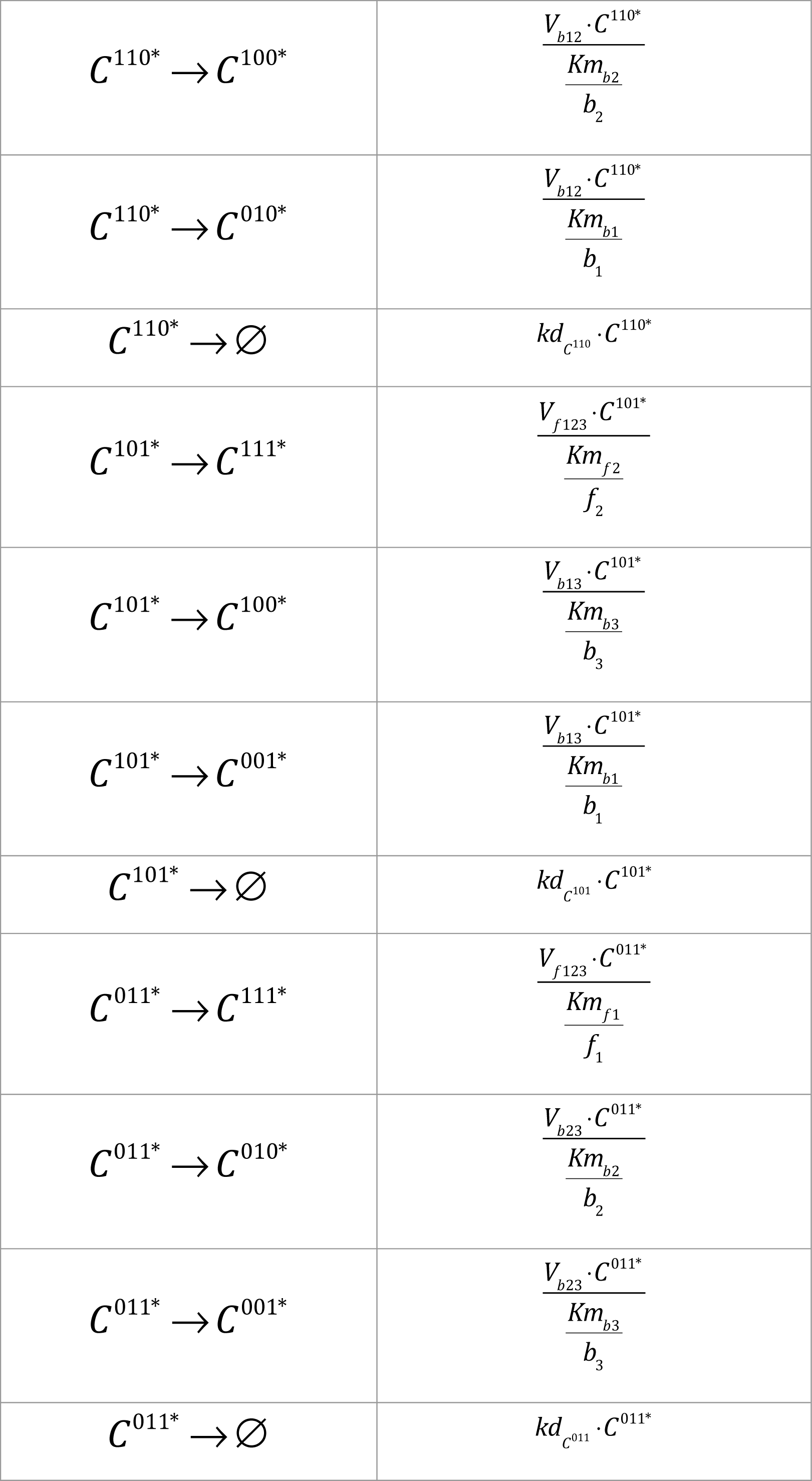
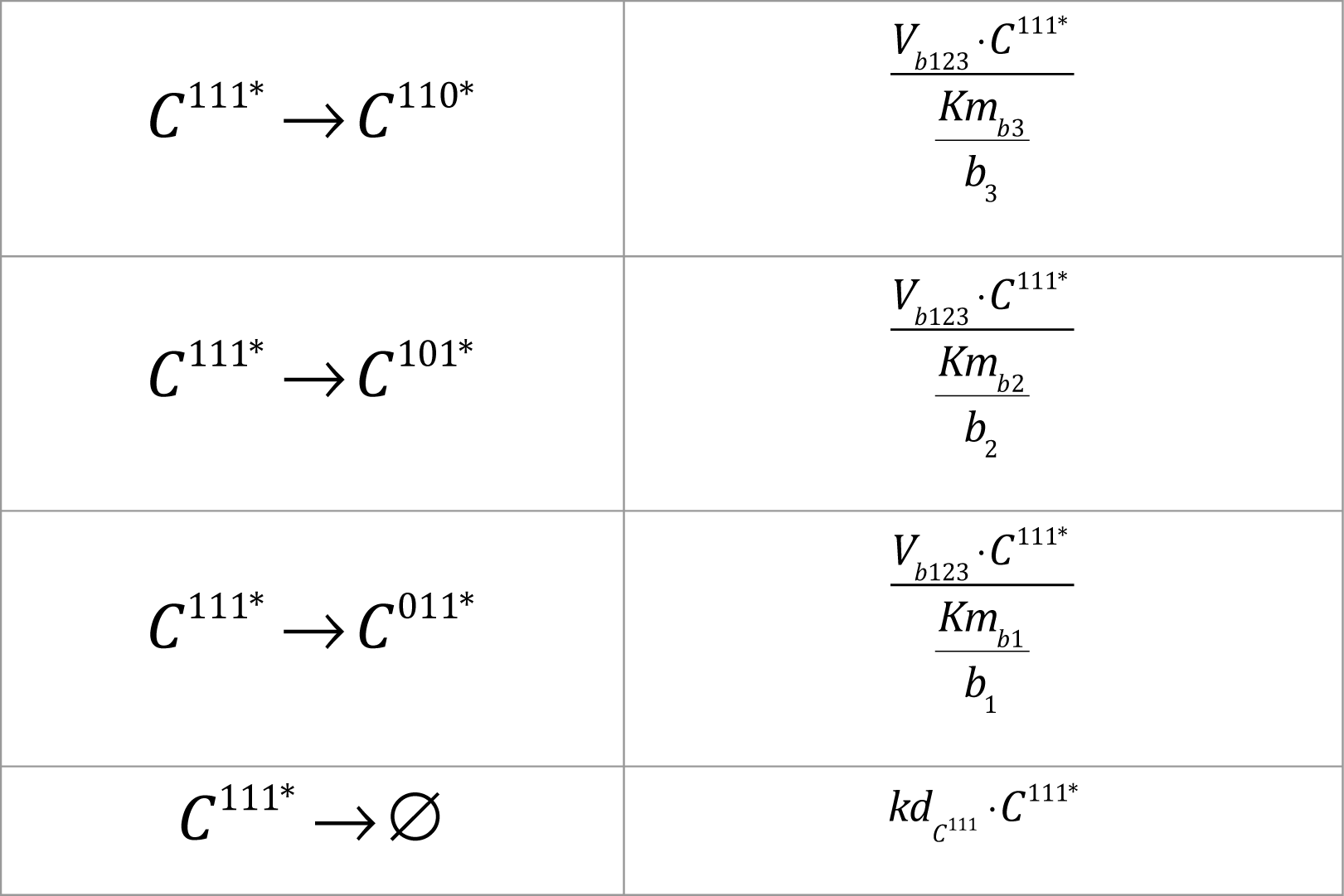
Propensity function used for stochastic simulations. In the column of the table, each line describes a reaction of the model. To each associated a rate, in the second column, that describes the probability of the reaction to happen at each time step of the stochastic simulation.

Where the competition terms *f* and *b* in S1 Table are defined as:

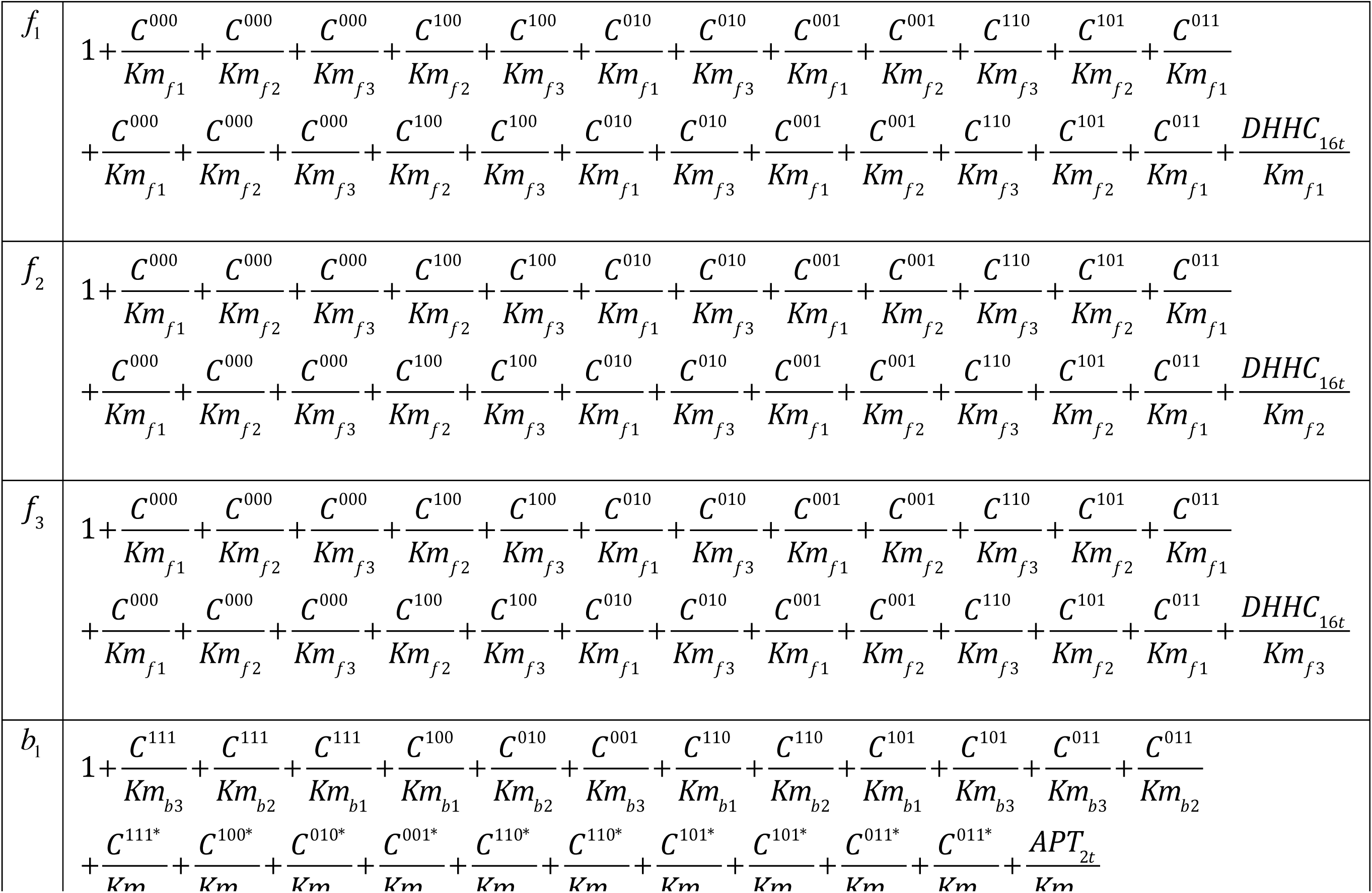
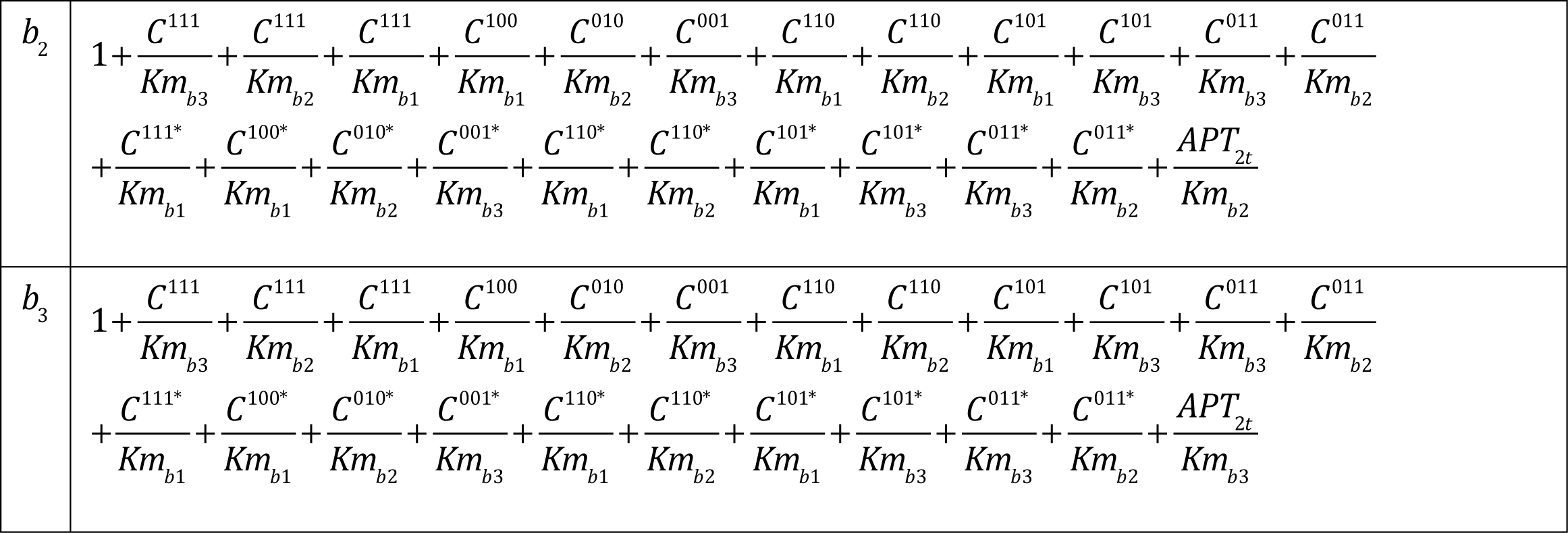

**Table S9.** Results of stochastic simulation when APT2 is silenced. The shows the average number of passages of a DHHC6 molecule in the different palmitoylation states when APT2 is silenced. The time spent in each state reported.

| siRNA APT |  |  |  |
| --- | --- | --- | --- |
| Species | Aver. Numb. passages /molecule | Median of the time spent per passage (h) | Median cumulative time (h) |
| C<sup>000</sup> | 1 | 0.9 | 0.9 |
| C<sup>100</sup> | 1 | 1.2 | 1.2 |
| C<sup>010</sup> | <0.1 | 0.7 | 0.7 |
| C<sup>001</sup> | <0.1 | <0.1 | <0.1 |
| C<sup>110</sup> | 0.5 | 2.7 | 2.7 |
| C<sup>101</sup> | 0.2 | 0.2 | 0.2 |
| C<sup>011</sup> | <0.1 | 9.9 | 9.9 |
| C<sup>111</sup> | 0.5 | 0.3 | 0.3 |

